# XRCC1 Enables the Efficient Local Search for DNA Damage by DNA Polymerase Beta

**DOI:** 10.64898/2026.08.24.746741

**Authors:** Spencer H. Thompson, Kaitlin M. DeHart, Matthew A. Schaich, Bret D. Freudenthal

**Affiliations:** Department of Biochemistry and Molecular Biology, University of Kansas Medical Center, Kansas City, KS 66160, USA; Department of Pharmacology and Chemical Biology, University of Pittsburgh School of Medicine, Pittsburgh, PA, USA; Molecular Biophysics and Structural Biology Program, University of Pittsburgh School of Medicine, Pittsburgh, PA, USA; Department of Cancer Biology, University of Kansas Medical Center, Kansas City, KS 66160, USA

**Keywords:** DNA damage and repair, base excision repair (BER), DNA polymerase β, x-ray repair cross-complementing 1, apurinic/apyrimidinic endonuclease 1 (APE1), single-molecule biophysics, optical tweezers, protein–DNA interactions

## Abstract

Oxidative DNA damage is a common threat to genomic integrity, arising from endogenous metabolic processes and environmental exposures. If unrepaired, such oxidative DNA damage promotes mutagenesis and genomic instability. Cells counter this through base excision repair (BER), a multi-step pathway requiring the coordinated action of several proteins. Central to BER, DNA polymerase beta (pol ꞵ) locates single-nucleotide (1-nt) gaps and inserts the correct nucleotide, while x-ray repair cross-complementing 1 (XRCC1) is a scaffold protein that forms a stable complex with pol ꞵ to coordinate BER factors at DNA damage. XRCC1 enhances BER efficiency, though the mechanism by which this occurs is unclear. Pol β is proposed to be recruited to DNA damage by undamaged DNA scanning interactions, but this behavior has not yet been directly observed. Additionally, the influence of other BER proteins on pol ꞵ recruitment, particularly XRCC1, remains unclear. Here, we used correlative optical tweezers-fluorescence microscopy to visualize DNA search and damage recognition by pol ꞵ and XRCC1. We characterize each factor individually, examine their behavior as the pol ꞵ-XRCC1 complex, and assess their interplay with apurinic/apyrimidinic endonuclease 1 (APE1), the enzyme upstream of pol ꞵ in BER. We find that pol ꞵ locates damage through 3D-diffusion, whereas XRCC1 exhibits both 3D- and 1D-diffusion. In combination, XRCC1 dramatically shifts pol β search towards 1D-diffusion, enabling interrogation of non-damaged DNA using both search mechanisms. When both APE1 and pol ꞵ are present, the pol ꞵ-1nt gap complex is highly stable, with APE1 largely unable to disrupt the damage-bound pol ꞵ. Together, these findings demonstrate that XRCC1 reshapes pol β search behavior to promote efficient local damage recognition, providing a mechanistic basis for how BER factors coordinate lesion detection and processing to maintain genomic stability.

**Significance Statement:** DNA repair proteins must locate rare sites of damage hidden within millions of undamaged bases. Using single-molecule imaging with optical tweezers, we directly visualized how DNA polymerase ꞵ and its scaffold partner XRCC1 search for and engage DNA damage. Alone, pol β finds damage exclusively through 3D collisions, whereas XRCC1 scans along DNA by 1D hopping. When the two proteins form a complex, XRCC1 confers its scanning ability on pol β, expanding the search strategies available for damage detection. These findings reveal a mechanism by which scaffold proteins remodel the damage search process of their partners, providing insight into how base excision repair is coordinated to maintain genome stability.

## Introduction

Reactive oxygen species (ROS) are a product of both environmental exposures and endogenous metabolic processes that represent an unavoidable cause of DNA damage(1–4). The primary pathway responsible for repairing ROS-induced nucleobase damage is base excision repair (BER)(3, 4). BER is initiated when damaged bases are excised by a DNA glycosylase and the resulting abasic sites are incised by apurinic/apyrimidinic endonuclease 1 (APE1), generating a single-strand break (Fig. 1A)(5–7). Subsequently, DNA polymerase beta (pol ꞵ) removes the remaining incised lesion and inserts the correct nucleotide at the one-nucleotide (1-nt) gap, generating a 3′ nick that is then sealed by ligase III alpha (LigIIIα)(8–11). During these final two enzymatic steps, pol ꞵ and LigIIIα interact with the scaffold protein x-ray repair cross-complementing 1 (XRCC1). XRCC1 can also be recruited to sites of DNA damage through its interaction with poly(ADP-ribose) chains, generated by poly(ADP-ribose) polymerase 1 (PARP1) recognition of strand breaks to facilitate assembly of downstream BER factors(12, 13). Failure to complete these final steps in BER can result in the persistence of single-strand DNA breaks (SSBs)(14–16).

**Figure 1.**
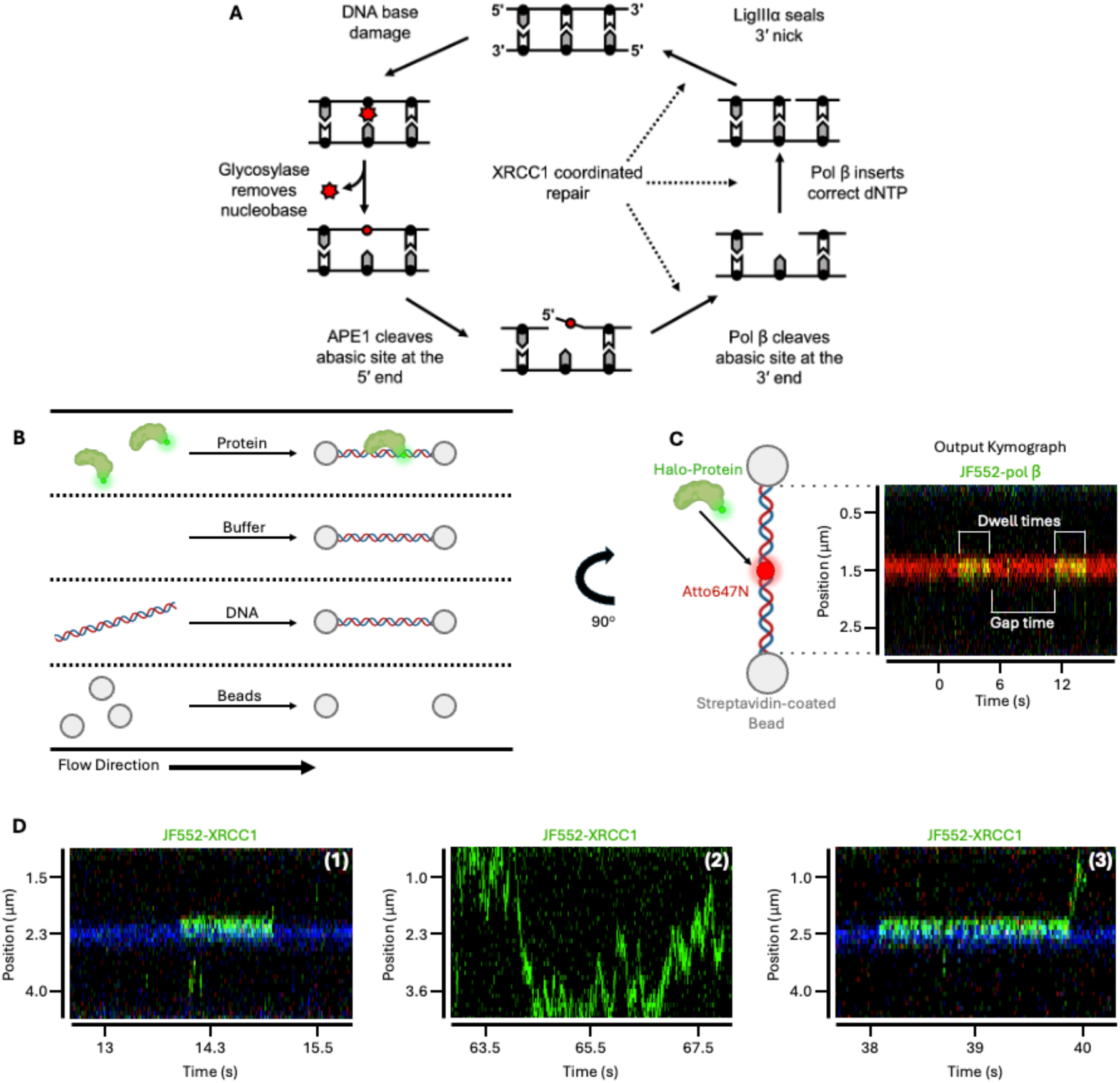
Experimental set-up with example kymograph analysis. (A) Base excision repair cycle. Reprinted and modified with permission from Whitaker, A.M. & Freudenthal, B.D. APE1: A skilled nucleic acid surgeon. DNA Repair (Amst) **71**, 93-100 (2018)(27) (B) Diagram of microfluidics flow cell showing four different chambers separated by laminar flow (dashed lines), each containing either streptavidin-coated polystyrene beads, biotinylated DNA containing sequence insert, buffer alone to check for the presence of a single DNA, or fluorescently labeled protein. A typical experiment set-up begins by capturing two beads and moving up through each chamber in subsequent steps. (C) Generation of a single kymograph showing a green Halo-protein binding the DNA near the site of the internal Atto647N fluorophore. Binding/unbinding events are used to calculate the protein dwell times and gap times. (D) Example kymographs demonstrating stationary (1), motile (2), and mixed protein binding (3). Green fluorescence represents JF552-labeled protein and blue fluorescence represents an internal Atto488 fluorophore at the 1-nt gap damage site. Made with Biorender.com

As a scaffold protein in BER, XRCC1 coordinates the activity of multiple repair factors including APE1, OGG1, LigIIIα and pol ꞵ(6, 17–19). The mechanisms by which XRCC1 modulates each of these interactions have been only partially defined. For example, work using cellular extracts has established that XRCC1 binds DNA nicks in complex with LigIIIα primarily through a direct, non-motile diffusion mechanism, but that the presence of XRCC1 is not required for LigIIIα binding at these sites (20). Whether XRCC1 engagement with pol ꞵ follows a similar pattern of direct, non-motile recruitment to damage sites, remains unknown. Notably, XRCC1 forms an exceptionally tight complex with pol ꞵ (*K*_D_≈10 nM), and several studies suggest that XRCC1 enhances the recruitment of pol ꞵ to sites of DNA damage(21–23). Recent cellular work indicates that the XRCC1-pol β interaction is required to suppress excessive PARP1 engagement at repair intermediates and thereby maintain rapid rates of BER(24). Interestingly, the pol ꞵ-XRCC1 interaction does not appear to be strictly required for pol ꞵ-mediated DNA repair *in vivo*, making the mechanism by which XRCC1 promotes efficient BER when bound to pol ꞵ unclear(25, 26). Additionally, the mechanism by which XRCC1 promotes pol ꞵ localization to DNA damage and, critically, whether XRCC1’s actively facilitates pol ꞵ damage search and thereby improves BER efficiency remains an open question.

To address how XRCC1 influences pol ꞵ damage search, and how BER proteins collectively navigate DNA repair intermediates, we used correlated optical tweezers and fluorescence microscopy (CTFM) to directly visualize fluorescently-tagged pol ꞵ and XRCC1 as they search for and bind DNA damage in real time. Our results indicate that XRCC1 primarily engages in 1D-diffusion along non-damaged DNA to facilitate its search for damage sites. In contrast, pol ꞵ almost exclusively binds to damage directly from bulk solution (3D-diffusion). Upon complex formation, the pol ꞵ-XRCC1 complex exhibits search behavior resembling XRCC1 alone, utilizing both 1D- and 3D-diffusion to locate DNA damage. In this complex, XRCC1 largely governs the 1D-diffusion behavior, while pol ꞵ primarily contributes to damage recognition and binding stability. In the presence of APE1, we find motile BER proteins bypass occupied damage sites rather than disrupting existing protein-DNA complexes. Together, these findings provide insight into how BER proteins concurrently search for and recognize DNA damage and reveal a mechanism by which XRCC1 alters pol ꞵ search behavior.

## Results

To understand how pol ꞵ and XRCC1 locate DNA damage, we used a CTFM experimental approach that enables direct visualization of recombinant pol ꞵ with an N-terminal HaloTag (Halo-pol ꞵ) and recombinant XRCC1 with a C-terminal HaloTag (XRCC1-Halo)(28). The Halo-pol ꞵ and XRCC1-Halo were labeled with JaneliaFluor Halo-Tag ligand 552 (JF552) and 635 (JF635). We generated a ∼12.6 kbp DNA substrate that was biotinylated at each end and contained either non-damaged DNA, a single-nucleotide (1-nt) gap, or 3′ phosphorylated nick (the substrate and product of pol ꞵ nucleotide insertion), with the damage site marked by an internal Atto647N or Atto488 fluorophore. Dual optical traps in the C-Trap were used to capture streptavidin-coated polystyrene beads and subsequently load a single DNA substrate between the beads to visualize protein-DNA interactions (Fig. 1B-C). The resulting kymographs were used to track the position of these proteins on the DNA over time, allowing us to determine binding lifetimes and diffusivity (Fig. 1C). From these trajectories, each binding event can be classified into one of three behaviors: stationary binding, in which the protein engages the DNA directly from solution and remains at a fixed position (3D diffusion); motile binding, in which the protein translocates along the DNA (1D diffusion); or mixed binding, which contains both components within a single event (Fig. 1D). This classification is used throughout to describe how each protein locates DNA damage.

### Pol ꞵ locates single-nucleotide gaps and 3**′** nicks using three-dimensional diffusion

To understand the modes of diffusion used by pol β during DNA damage search and recognition, we studied pol β binding to undamaged DNA, 1-nt gap (substrate) and 3′ nick (product) containing DNA. Our 1-nt gap experiments were conducted in the absence of dNTPs to prevent nucleotide addition which would complicate mechanistic interpretation. In the presence of either the 1-nt gap or 3′ nick, pol ꞵ demonstrated exclusively stationary behavior at the damage site (binding directly from solution), indicative of protein binding through 3D-diffusion (Fig. 2A-C, Supplementary Fig. S1-2). Notably, this is the only behavior we observe pol ꞵ using on 1-nt gaps and 3′ nicks, over the >1000 observed events, to locate these damage sites, without observing the protein perform motile 1D-diffusion. However, given the spatial resolution of our assay, we cannot necessarily exclude short-range 1D diffusion (on the order of 200–300 bp) occurring prior to or immediately surrounding pol ꞵ damage recognition, as such movements would fall below our limit of detection. To confirm that pol ꞵ primarily uses 3D-diffusion for DNA search, we also performed experiments on non-damaged DNA (Fig. 2A). With this substrate, we did not observe pol β binding events on our non-damaged substrate despite testing multiple protein concentrations (up to 1.5 nM) and utilizing faster line times (10 ms). The inability to observe binding events under these conditions suggests pol ꞵ has a weak affinity for undamaged DNA or probes the non-damaged DNA with dwell times that are shorter than the temporal resolution of the C-Trap. This is also consistent with the binding affinity of pol ꞵ to non-damaged DNA being poor (500-700 µM)(29).

**Figure 2.**
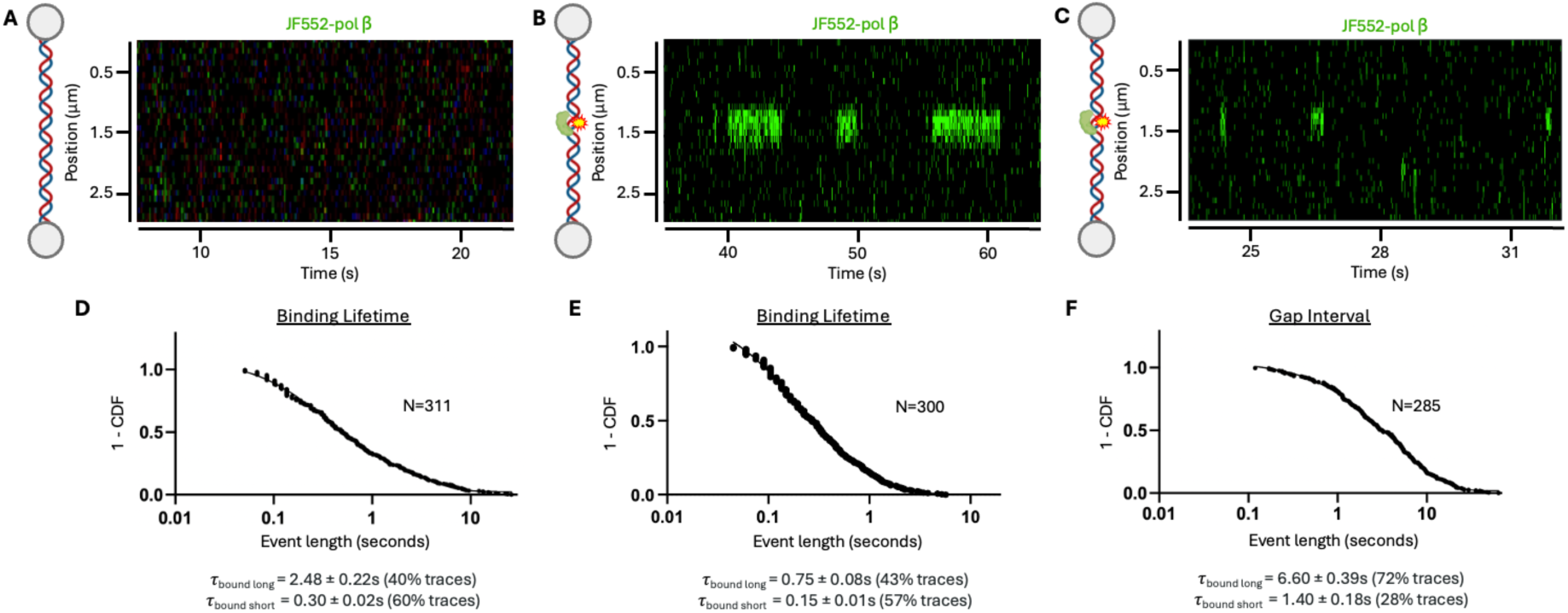
Pol ꞵ is nonmotile and uses 3D-diffusion to locate DNA damage. (A) Example kymograph of pol ꞵ and non-damaged DNA demonstrating a lack of pol ꞵ binding events. (B) Example kymograph of nonmotile pol ꞵ binding directly to a one-nucleotide gap. (C) Example kymograph of nonmotile pol ꞵ binding directly to a 3′ nick. (D) Cumulative distribution function (CDF) plot of pol ꞵ dwell times on DNA containing a single-nucleotide gap stretched to 10 pN. Dwell times of short and long-lived pol ꞵ binding populations are indicated. (E) CDF plot of pol ꞵ dwell times on DNA containing a 3′ nick stretched to 10 pN. (F) CDF plot of pol ꞵ gap times on DNA containing a single-nucleotide gap stretched to 10 pN.

To determine how stably pol β engages a 1-nt gap during DNA damage recognition, we quantified the duration of individual pol ꞵ binding events at 1-nt gaps (dwell times, Fig. 1C) and from these determined a representative binding lifetime (τ). Notably, two dwell time populations emerged, possibly reflecting different binding conformations of pol ꞵ at the damage site: one population with a shorter binding lifetime (hereafter, short-lived population) and one with a longer binding lifetime (hereafter, long-lived population). At 10 pN, the short-lived population represented 60% of events (average binding lifetime of 0.30 ± 0.02s), whereas the long-lived population represented 40% of events (average binding lifetime of 2.48 ± 0.22s), giving an average weighted binding lifetime of 1.17 ± 0.14s at the 1-nt gap (Fig. 2D, Table 1). In comparison, the populations of pol ꞵ binding at the 3′ nick demonstrated a short-lived population representing 57% of events (average binding lifetime of 0.15 ± 0.01s), whereas the long-lived population represented 43% of events (average binding lifetime of 0.75 ± 0.08s), giving an average weighted binding lifetime of 0.41 ± 0.04s, shorter than at the 1-nt gap and consistent with a comparatively weaker interaction of pol ꞵ at the product of nucleotide insertion (Fig. 2E, Table 1). These results demonstrate that pol ꞵ exhibits greater ability to form a long-lived binding complex at its substrate (1-nt gap) than its product (3′ nick), suggesting that nucleotide-binding competent conformations are preferentially stabilized at the gap.

**Table 1.** Dwell and gap time intervals for pol ꞵ at 10-40 pN at 1-nt gaps or 3′ nicks. Dwell times correspond to stationary pol ꞵ binding events at each damage site, whereas gap times represent the interval between successive pol ꞵ binding events. When fit to exponential decay functions, several dwell/gap datasets were best described by multiple kinetic populations. These populations are reported as *τ*_bound short_, *τ*_bound medium_, or *τ*_bound long_, along with the percentage of events assigned to each population. Weighted average lifetimes were calculated from these values and used to determine the mean dwell or gap lifetime, and corresponding binding affinity.

|  | One-nucleotide gaps |  |  | 3' nicks |  |  |
| --- | --- | --- | --- | --- | --- | --- |
| Dwell Times |  |  |  |  |  |  |
|  | 10pN | 20pN | 40pN | 10pN | 20pN | 40pN |
| $\tau_{\text{bound short}}$ | 0.30 ± 0.02s | 0.10 ± 0.01s | 0.07 ± 0.004s | 0.15 ± 0.01s | 0.21 ± 0.02s | 0.17 ± 0.01s |
| $\tau_{\text{bound medium}}$ | - | - | - | - | - | - |
| $\tau_{\text{bound long}}$ | 2.48 ± 0.22s | 0.96 ± 0.20s | 1.33 ± 0.92s | 0.75 ± 0.08s | 0.94 ± 0.06s | 1.11 ± 0.15s |
| N | 311 | 329 | 241 | 300 | 306 | 388 |
| Distribution | 60% short<br>40% long | 80% short<br>20% long | 90% short<br>10% long | 57% short<br>43% long | 34% short<br>66% long | 74% short<br>26% long |
| $\tau_{\text{average}}$ | 1.17 ± 0.14 | 0.27 ± 0.04s | 0.19 ± 0.09s | 0.41 ± 0.04s | 0.69 ± 0.04s | 0.41 ± 0.04s |
| Gap Times |  |  |  |  |  |  |
|  | 10pN | 20pN | 40pN | 10pN | 20pN | 40pN |
| $\tau_{\text{bound short}}$ | 1.40 ± 0.18s | - | 0.99 ± 0.05s | 2.48 ± 0.09s | 0.52 ± 0.09s | 2.79 ± 0.08s |
| $\tau_{\text{bound medium}}$ | - | 5.66 ± 0.07s | 5.28 ± 0.28s | - | 2.13 ± 0.19s | - |
| $\tau_{\text{bound long}}$ | 6.60 ± 0.39s | - | 24.01 ± 1.70s | 11.88 ± 2.26s | 8.00 ± 0.82s | 13.31 ± 3.32s |
| N | 285 | 313 | 225 | 311 | 289 | 360 |
| Distribution | 28% short<br>72% long | 100% | 20% short<br>45% medium<br>35% long | 75% short<br>25% long | 11% short<br>57% medium<br>32% long | 84% short<br>16% long |
| $\tau_{\text{average}}$ | 5.14 ± 0.29s | 5.66 ± 0.07s | 10.98 ± 0.61s | 4.83 ± 0.57s | 3.83 ± 0.29s | 4.47 ± 0.53s |
| $K_D$ | 2.06 ± 0.35nM | 10.5 ± 1.6nM | 28.9 ± 13.9nM | 5.9 ± 0.9nM | 2.78 ± 0.29nM | 5.45 ± 0.83nM |

To investigate the effect of relevant mechanical forces on pol ꞵ binding, we next collected binding events at 20 pN and 40 pN applied along the DNA, accomplished by extending the DNA held under tension by the dual optical traps. Cellular processes such as transcription can generate forces on the order of ∼30 pN during RNA polymerase-driven elongation, which can alter the binding behavior of DNA-interacting proteins including their on-rates(28, 30–32). This stretching may compete with the ∼90° DNA bend that pol ꞵ induces upon gap binding, a conformational change required for catalysis, and may promote gap widening or base fraying that destabilizes pol ꞵ engagement(33). At 20 pN, the short-lived population at the 1-nt gap increased from 60% (at 10 pN) to 80% of events (0.10 ± 0.01s), with the long-lived population reduced to 20% (0.96 ± 0.20s), giving an average weighted binding lifetime of 0.27 ± 0.04s at 20 pN. At 40 pN, 90% of events belonged to the short-lived population (0.07 ± 0.004s) and 10% to the long-lived population (1.33 ± 0.92s), giving an average weighted binding lifetime of 0.19 ± 0.09s (Table 1). Cumulatively, increasing force on the DNA shifted events toward the short-lived population and reduced overall binding lifetimes, indicating that tension destabilizes pol ꞵ retention at 1-nt gaps and suppresses the formation of stable long-lived complexes. This force-dependent effect may suggest distinct modes of DNA engagement, with short-lived interactions potentially corresponding to transient sampling states and long-lived interactions representing more stably engaged conformations at the damage site. In contrast, pol ꞵ binding lifetimes at 3′ nicks were not notably altered by force (0.41 ± 0.04s at 10 pN, 0.69 ± 0.04s at 20 pN, and 0.41 ± 0.04s at 40 pN), suggesting that the 1-nt gap is uniquely susceptible to force-dependent structural perturbations compared to the 3′ nicks (Table 1). This may be the result of pol β adopting a binding mode on the nick that does not require the same degree of DNA bending as at the 1-nt gap, potentially reflecting engagement of the lyase domain with the 5′ phosphate in a partially bent intermediate state similar to those observed in nucleosome-bound structures where the lyase domain is bound while the polymerase domain remains disengaged(34).

To further characterize the kinetics of pol β binding and its ability to re-engage DNA lesions after dissociation, we determined the gap times (intervals between successive binding events) at the 1-nt gap under each applied force (Fig. 2F). Together with dwell times, gap times allow us to calculate on- and off-rates, and thus the equilibrium dissociation constant (K_D_) for pol ꞵ at each damage substrate. Full gap time distributions at each force are reported in Table 1. Using the average weighted binding lifetime and gap interval, the K_D_ for pol ꞵ for 1-nt gaps was 2.06 ± 0.35 nM at 10 pN, 10.5 ± 1.6 nM at 20 pN, and 28.9 ± 13.9 nM at 40 pN, suggesting that pol ꞵ binding is sensitive to gap size, consistent with prior reports(35). This force-dependent decrease in binding affinity was driven by shorter dwell times and extended gap times, possibly reflecting force-induced structural changes such as gap widening or base fraying that impair pol ꞵ binding. Interestingly, no analogous force-dependence was observed for pol ꞵ at 3′ nicks, where the K_D_ was 5.9 ± 0.9 nM at 10 pN, 2.8 ± 0.3 nM at 20 pN, and 5.5 ± 0.8 nM at 40 pN.

### XRCC1 locates single-nucleotide gaps and 3**′** nicks using one-dimensional and three-dimensional diffusion

To understand how XRCC1 searches non-damaged DNA, we performed single molecule imaging of XRCC1 on our non-damaged DNA substrate (Fig. 3A). We found that XRCC1 exhibits predominantly motile behavior (93% of events) with few exhibiting stationary binding (6% of events) or a mix of both motile and stationary binding (1% of events) (Fig. 3B). To our knowledge, this is the first reporting that XRCC1 can conduct 1D-diffusion along a non-damaged DNA substrate, demonstrating that a BER scaffold protein can actively patrol DNA in search of damage. To understand the timescale of XRCC1-mediated damage search, we next quantified the duration of XRCC1 search on non-damaged DNA. We identified two populations of XRCC1 binding events, a short-lived population representing 79% of events where XRCC1 bound for 0.12 ± 0.01s and a long-lived population representing 21% of events bound for 0.93 ± 0.20s (Fig. 3C, Table 2). This represents an average weighted binding lifetime of 0.29 ± 0.04s, the approximate length of time XRCC1 alone conducts 1D damage search before dissociating from DNA.

**Figure 3.**
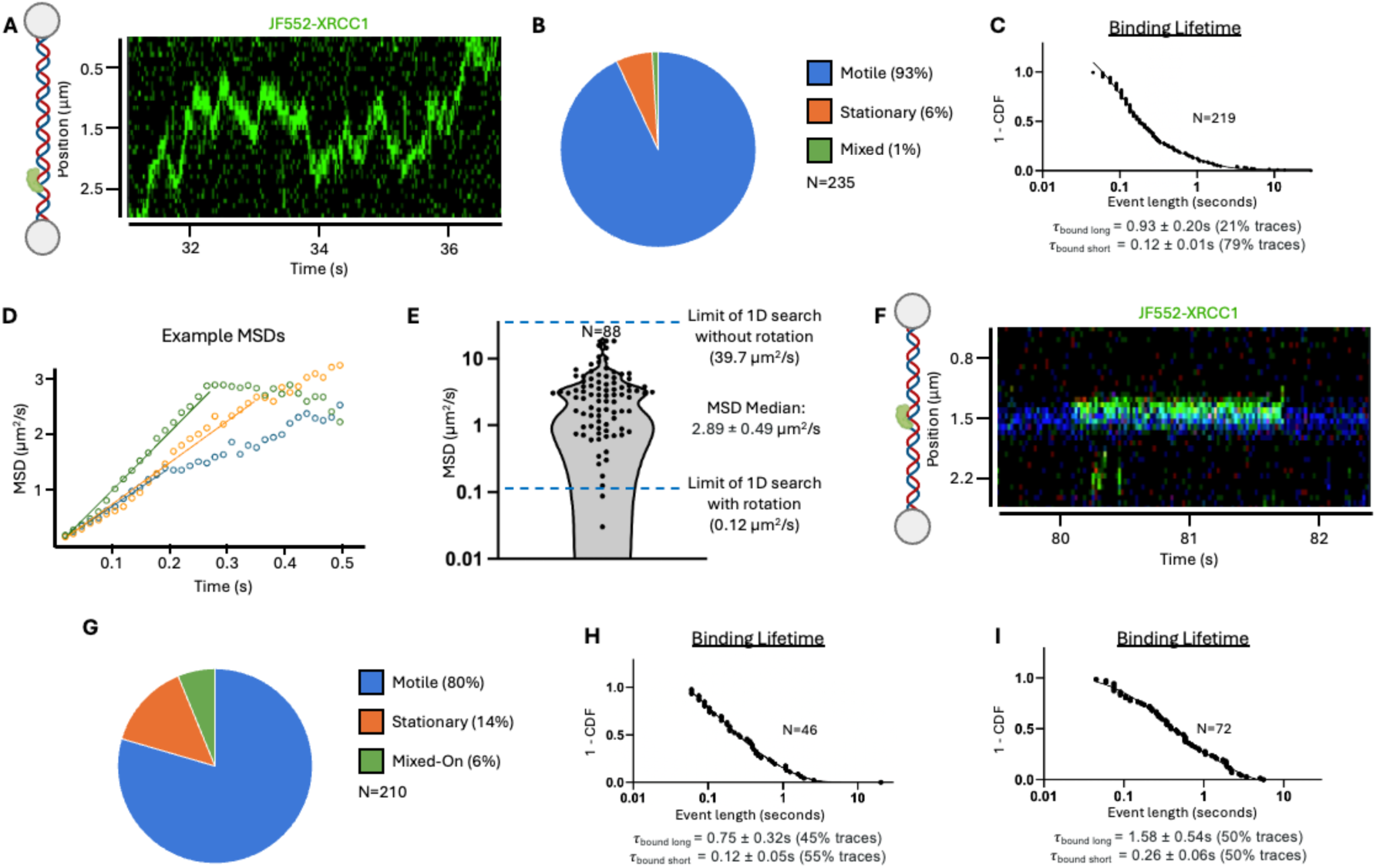
XRCC1 is motile and uses 1D- & 3D-diffusion to locate DNA damage. (A) Example kymograph of XRCC1 searching through the length of the ∼12.6kbp non-damaged DNA substrate. (B) Percentage of XRCC1 events on non-damaged DNA that were either motile, stationary, or mixed (elements of both motile and stationary behavior). (C) Cumulative distribution function (CDF) plot of the XRCC1 dwell times on non-damaged DNA. (D) Example MSD plots denoting the linear portion of the MSD versus time that was fit to determine a diffusion coefficient from a single binding event. (E) Violin plot of the diffusion coefficients derived from MSD analysis of XRCC1 on non-damaged DNA. (Median ± SEM*√(π/2)) (F) Example kymograph of nonmotile XRCC1 binding directly to a one-nucleotide gap. (G) Percentage of XRCC1 events on DNA containing a one-nucleotide gap that were motile, stationary, or mixed. (H) CDF plot of the XRCC1 dwell times on DNA containing a 1-nt gap. (I) CDF plot of the XRCC1 dwell times on DNA containing a 3′ nick.

**Table 2.** Dwell time intervals for XRCC1 and the pol ꞵ-XRCC1 complex on non-damaged DNA, 1-nt gaps or 3′ nicks. (A) Dwell times correspond to stationary XRCC1 or pol ꞵ-XRCC1 binding events at each damage site. When fit to exponential decay functions, several dwell/gap datasets were best described by multiple kinetic populations. These populations are reported as *τ*_bound short_ or *τ*_bound long_, along with the percentage of events assigned to each population. Weighted average lifetimes were calculated from these values.

|  | Non-damaged |  | One-nucleotide gaps |  | 3' nicks |  |
| --- | --- | --- | --- | --- | --- | --- |
| Dwell Times |  |  |  |  |  |  |
|  | XRCC1 | Pol β-XRCC1 | XRCC1 | Pol β-XRCC1 | XRCC1 | Pol β-XRCC1 |
| $\tau_{\text{bound short}}$ | 0.12 ± 0.01s | 0.10 ± 0.01s | 0.12 ± 0.05s | 0.49 ± 0.07s | 0.26 ± 0.06s | 0.14 ± 0.10s |
| $\tau_{\text{bound long}}$ | 0.93 ± 0.20s | 0.87 ± 0.30s | 0.75 ± 0.32s | 2.12 ± 0.89s | 1.58 ± 0.54s | 1.14 ± 0.21s |
| N | 219 | 257 | 46 | 152 | 72 | 66 |
| Distribution | 79% short<br>21% long | 81% short<br>19% long | 55% short<br>45% long | 59% short<br>41% long | 50% short<br>50% long | 21% short<br>79% long |
| $\tau_{\text{average}}$ | 0.29 ± 0.04s | 0.25 ± 0.06s | 0.40 ± 0.15s | 1.16 ± 0.37s | 0.92 ± 0.27s | 0.93 ± 0.17s |

We quantified the diffusivity of XRCC1 on non-damaged DNA to determine how quickly the protein undergoes 1D-diffusion and the mechanism of 1D-diffusion that is being employed. To do this, we calculated the mean square displacement (MSD) of individual motile binding events to obtain diffusion coefficients and subsequently the median diffusion coefficient (D_med_) from the total population of motile events, which represent 93% of the total events observed with XRCC1 on non-damaged DNA (Fig. 3D). We determined the median diffusivity of XRCC1 was 2.9 ± 0.5 µm^2^/s. This means that if a single XRCC1 is bound for the length of our average weighted binding lifetime on non-damaged DNA (0.29 ± 0.04s), then based on the median diffusivity of XRCC1, the protein can scan 3.8 ± 0.4 kbp of DNA in a single binding event. The 1D diffusivity can be further broken down into two modes of scanning, either movement that is rotationally-coupled to the structure of the DNA helix (referred to as “sliding”) or non-rotationally-coupled movement (referred to as “hopping”)(36). The maximum diffusivity of a protein on a single strand of DNA using 1D movement by either sliding or hopping is limited by the drag of the protein along the DNA using these movement mechanics. We estimated the maximum diffusion coefficient of XRCC1 when hopping is 39.7 µm^2^/s while the limit for sliding is 0.12 µm^2^/s (see Methods). Because our median diffusion coefficient of XRCC1 was significantly above the limit for sliding (with <2.5% of data below this threshold) yet below the limit for hopping, this indicates that XRCC1 undergoes hopping during 1D search (Fig. 3E). From these results, we show that XRCC1 can rapidly traverse along non-damaged DNA and primarily does so via hopping mechanisms.

Having characterized how XRCC1 searched through non-damaged DNA, we next observed if/how XRCC1 locates DNA damage using 1-nt gaps or 3′ nicks contained within the DNA. With the 1-nt gap present, we observed a mix of behavior where 80% of XRCC1 events were motile, 14% were stationary, and 6% exhibited a mix of both motile and stationary behavior (encompassing all binding events on the 1-nt gap substrate, regardless of whether they engaged the damage site) (Fig. 3F-G, Supplementary Fig. S3). Between the two substrates, XRCC1 bound slightly longer to the nick DNA (0.92 ± 0.27s) than the gap DNA (0.40 ± 0.15s), indicating preference for that substrate (Fig. 3H-I, Table 2). Of note, the presence of damage did not significantly alter the behavior of XRCC1 compared with non-damaged DNA, with most events still showing motile behavior. This suggests that XRCC1 alone can use both 3D-diffusion and 1D hopping to scan for DNA damage but does not have a strong specificity for damage versus non-damaged DNA.

### Formation of the pol ꞵ-XRCC1 complex alters search dynamics and damage affinity

To understand how the pol ꞵ-XRCC1 complex searches non-damaged DNA, we first observed this complex interacting with our non-damaged DNA substrate. Previously, we did not observe instances of pol β binding to non-damaged DNA in the absence of XRCC1. Here, with the addition of XRCC1, we observed JF635-pol β and JF552-XRCC1 dual-color colocalization on non-damaged DNA, indicating formation of the pol β-XRCC1 complex (Fig. 4A, Supplementary Fig. S4). We observed that under these conditions, pol ꞵ-XRCC1 exhibits predominantly motile behavior (84% of events) with some events undergoing stationary binding (12% of events) or a mix of both motile and stationary binding (4% of events) (Fig. 4B). Given the highly motile behavior of this complex, similar to XRCC1 alone, we hypothesize that XRCC1 is facilitating pol ꞵ DNA damage search using 1D-diffusion to scan through non-damaged DNA. Consistent with this, when we used the L301R/V303R/V306R pol ꞵ variant, previously shown not to interact with XRCC1, pol ꞵ was still able to bind DNA damage, but formation of the motile pol ꞵ-XRCC1 complex did not occur (Supplementary Fig. S5)(37, 38) Next, to understand how the pol ꞵ-XRCC1 complex locates 1-nt DNA gaps, we observed this complex searching for and binding at that damage site (Fig. 4C and Supplementary Fig. S4). We observed that on gapped DNA, the complex was motile on non-damaged DNA in 53% of events, underwent stationary binding at the gap 35% of the time, and exhibited mixed behavior at the gap 12% of the time (Fig. 4D). We also observed how this complex recognizes 3′ nicks. We found that on nicked DNA 39% of events were motile, 53% of events underwent stationary binding at the nick, and 8% showed mixed behavior binding the nick (Fig. 4E). Notably, in the presence of either a 1-nt gap or 3′ nick, the pol ꞵ-XRCC1 complex can identify damage via motile (1D-diffusion) search, a search mechanism not observed for pol ꞵ alone. These findings indicate that pol ꞵ and XRCC1 can form a motile complex that actively scans DNA to allow pol ꞵ search capacity beyond static, 3D sampling and allows the complex to identify DNA damage through both 1D- and 3D-diffusion.

**Figure 4.**
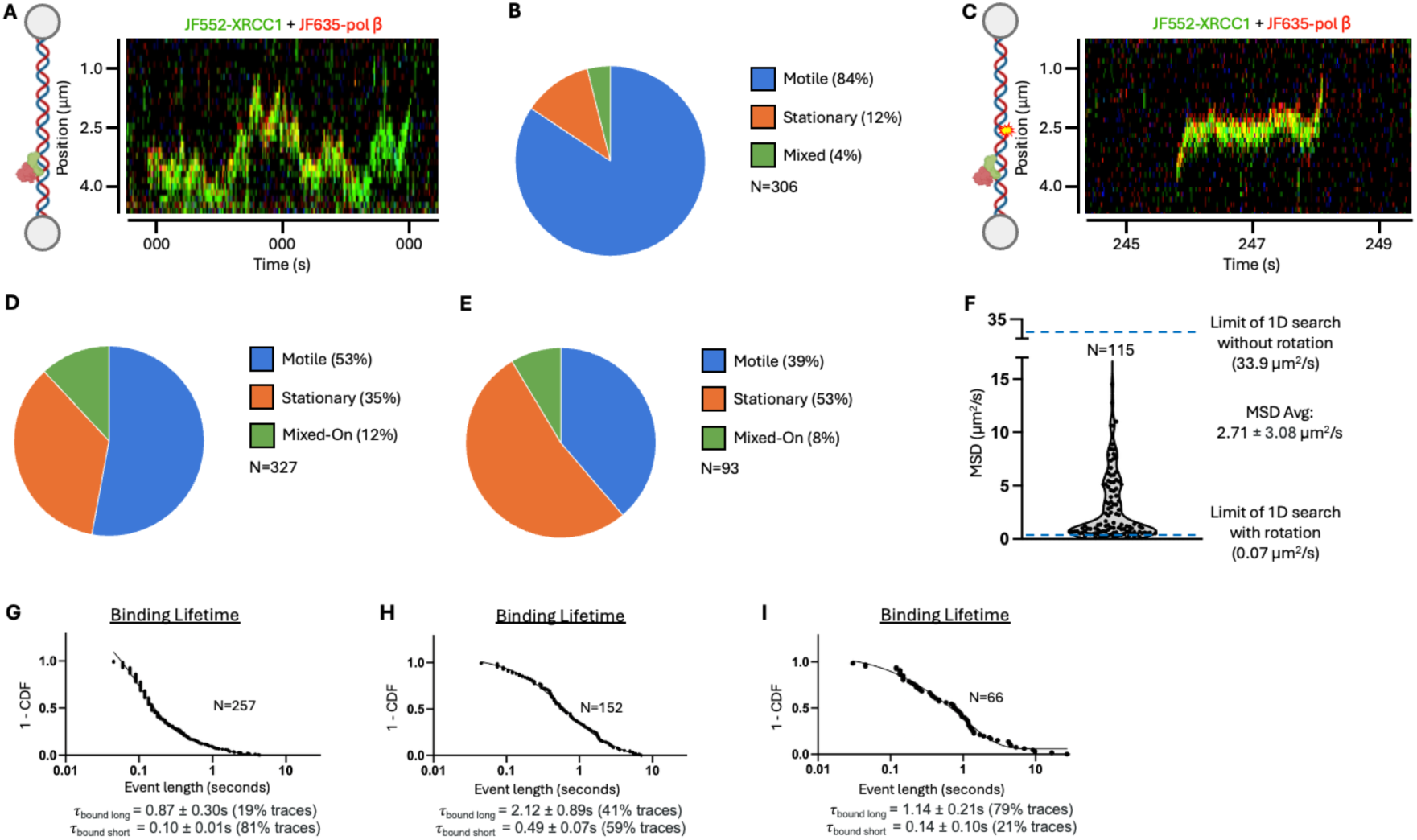
The pol ꞵ/XRCC1 complex uses both 1D- & 3D-diffusion to locate DNA damage. (A) Example kymograph of a motile pol ꞵ (red)-XRCC1 (green) complex searching through non-damaged DNA. (B) Percentage of pol ꞵ-XRCC1 complex events on non-damaged DNA that were either motile, stationary, or mixed. (C) Example kymograph of a mixed pol ꞵ-XRCC1 complex using 1D-diffusion to locate a one-nucleotide gap, sampling the damage site, and continuing 1D search beyond the damage. (D) Percentage of pol ꞵ-XRCC1 events on DNA containing a one-nucleotide gap that were motile, stationary, or mixed. (E) Percentage of pol ꞵ-XRCC1 events on DNA containing a 3′ nick that were motile, stationary, or mixed. (F) Violin plot of the diffusion coefficients derived from MSD analysis of pol ꞵ-XRCC1 on non-damaged DNA. (Median ± SEM*√(π/2)) (G) CDF plot of the pol ꞵ-XRCC1 complex dwell times on non-damaged DNA. (H) CDF plot of pol ꞵ-XRCC1 complex dwell times on DNA containing a 1-nt gap, defined as intervals during which both proteins are simultaneously present at the damage site. (I) CDF plot of the pol ꞵ-XRCC1 complex dwell times on DNA containing a 3′ nick.

We next quantified the binding lifetimes of this complex on non-damaged, 1-nt gap, or 3′ nick DNA. This revealed that on non-damaged DNA, motile pol ꞵ-XRCC1 has a short-lived population representing 81% of events where the complex was bound for 0.10 ± 0.01s and a long-lived population representing 19% of events where the complex was bound for 0.87 ± 0.30s (Fig. 4G, Table 2). This represents an average binding lifetime on non-damaged DNA of 0.25 ± 0.06s, which is within error of XRCC1 alone on this substrate (0.29 ± 0.04s). On DNA containing a 1-nt gap, we find the complex binds the gap, defined here as any interval in which both proteins were simultaneously present, with a short-lived population representing 59% of events where the complex was bound for 0.49 ± 0.07s and a long-lived population representing 41% of events where the complex was bound for 2.12 ± 0.89s (Fig. 4H, Table 2). This represents an average binding lifetime on DNA containing a 1-nt gap of 1.16 ± 0.37s, essentially the same as pol ꞵ alone (1.17 ± 0.14s, Table 1), indicating retention at the 1-nt gap is mainly driven by pol ꞵ. Finally, on DNA containing a 3′ nick, we find the complex binds the nick with a short-lived population representing 21% of events where the complex was bound for 0.14 ± 0.10s and a long-lived population representing 79% of events where it was bound for 1.14 ± 0.21s (Fig. 4I, Table 2). This represents an average binding lifetime on DNA containing a 3′ nick of 0.93 ± 0.17s, twofold longer than pol ꞵ alone (0.41 ± 0.04s) and similar to XRCC1 alone 0.92 ± 0.27s. The extended nick dwell time of the pol ꞵ-XRCC1 complex relative to pol ꞵ alone may reflect XRCC1-mediated stabilization at the nick, possibly pointing to the established role of XRCC1 interacting with DNA ligase IIIα at nick substrates. In contrast, XRCC1 does not appear to stabilize pol ꞵ at the 1-nt gap, demonstrated by the nearly identical dwell times of the pol ꞵ-XRCC1 complex compared to pol ꞵ alone.

We further determined the median diffusivity of pol ꞵ-XRCC1 on non-damaged DNA was 1.14 ± 0.36 µm^2^/s (Fig. 4F). If a single pol ꞵ-XRCC1 is bound for the length of our average weighted binding lifetime on non-damaged DNA (0.25 ± 0.06s), then based on the median diffusivity of pol ꞵ-XRCC1, the complex is capable of scanning 2.2 ± 0.4 kbp of DNA in a single binding event. This is slightly lower in diffusivity and average scanning distance than XRCC1 alone, possibly caused by drag and/or DNA damage probing by pol ꞵ. We estimated the maximum diffusion coefficient of the pol ꞵ-XRCC1 complex when hopping is 33.9 µm^2^/s while the limit for sliding is 0.07 µm^2^/s. Given our median diffusion coefficient of pol ꞵ-XRCC1 is 1.14 µm^2^/s, significantly above the limit for sliding yet below the limit for hopping, this indicates that pol ꞵ-XRCC1 mostly undergoes hopping. From these results, we show that pol ꞵ-XRCC1 can rapidly traverse along non-damaged DNA via 1D-diffusion, primarily using hopping during search, though we note this conclusion relies on theoretical estimates of diffusion limits (see Methods).

We next investigated how the pol ꞵ-XRCC1 complex assembles and disassembles at DNA damage. At 1-nt gaps, the most common mode of interaction was binding and dissociating as a preformed complex (59% of assembly events; 53% of disassembly events), with complex assembly at damage predominantly occurring via 3D-diffusion (82%) (Fig. 5A-B) rather than undergoing 1D-diffusion to locate the damage site (18%). When sequential binding was observed, pol ꞵ most frequently bound the 1-nt gap before XRCC1 (91%) and dissociated after XRCC1 (86%), indicating that pol ꞵ engages the lesion more stably than XRCC1. A similar pattern was observed at 3′ nicks. In that case, the complex binding and dissociating as a preformed complex (45% of assembly events; 42% of disassembly events) was still favored over any one protein binding first (Fig. 5C-D). Sequential binding at the nick typically involved pol ꞵ binding first (66%) and subsequently with XRCC1 dissociating first (68%). Together, these results reveal an asymmetry in complex behavior, where pol ꞵ appears to dominate lesion engagement and retention, while XRCC1 may contribute more to damage search than stable binding. This is consistent with a model in which XRCC1 contributes to DNA damage search within the complex but not necessarily its retention at all DNA lesions, which is primarily mediated by pol ꞵ(22).

**Figure 5.**
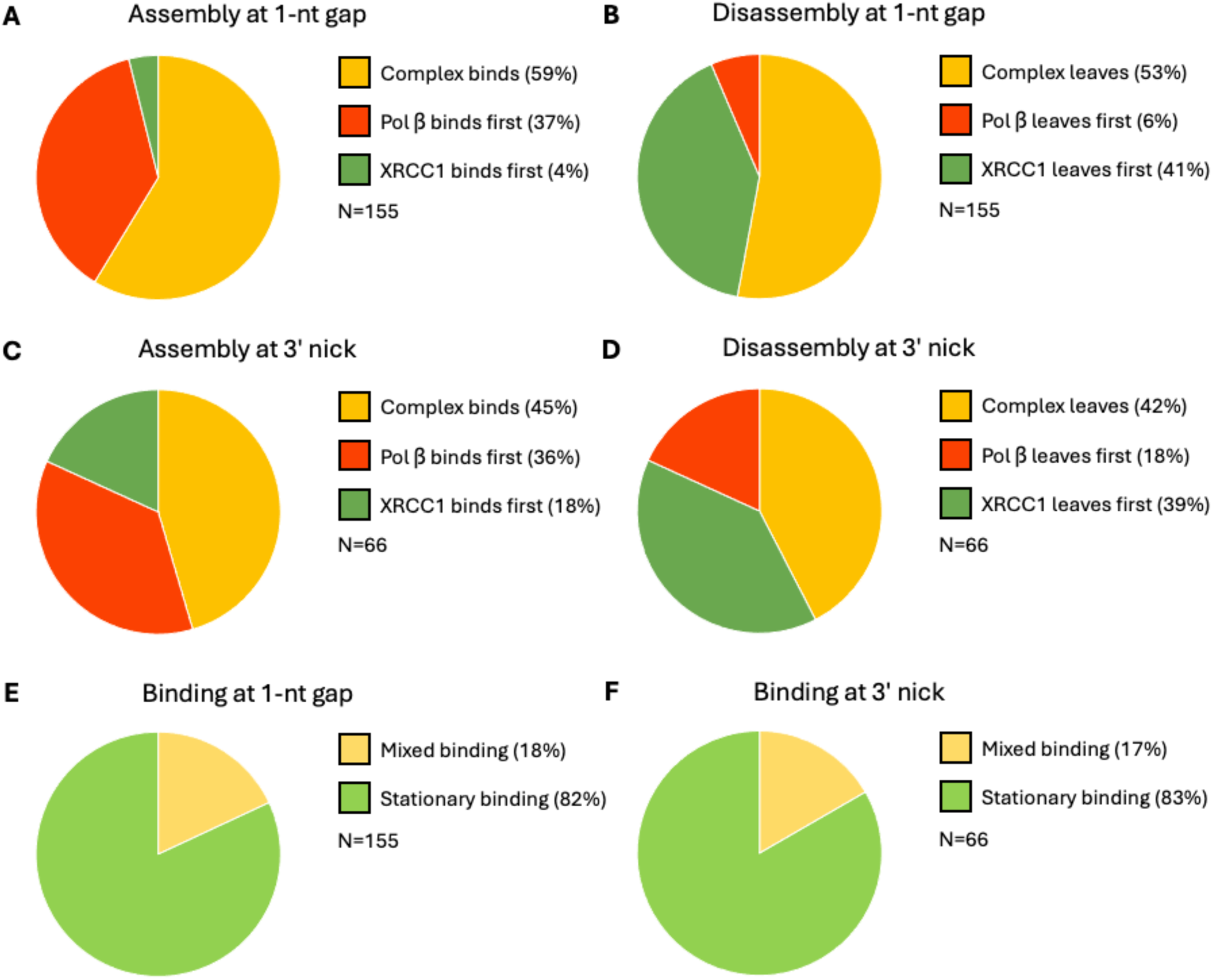
Assembly and disassembly order of the pol ꞵ-XRCC1 complex. (A) Percentage of assembly events where the pol ꞵ-XRCC1 complex bound to a 1-nt gap preassembled, with pol ꞵ binding before XRCC1, or with XRCC1 binding before pol ꞵ. (B) Percentage of disassembly events where the pol ꞵ-XRCC1 complex dissociated from the 1-nt gap as a complex, with pol ꞵ disassociated before XRCC1, or with XRCC1 disassociating before pol ꞵ. (C) Same as in A. but for DNA containing a 3′ nick. (D) Same as in B. but for DNA containing a 3′ nick. (E) Percentage of assembly events where the pol ꞵ-XRCC1 complex binds the 1-nt gap via 3D (stationary) binding or by 1D/3D (mixed) binding. (F) Same as in E. but for DNA containing a 3′ nick.

### Simultaneous damage search of pol ꞵ or XRCC1 with APE1

Previous work has shown that APE1 is capable of motile 1D-diffusion along non-damaged DNA(39, 40). It has further been demonstrated that interactions between APE1 and pol β can allow direct channeling of DNA repair intermediates between the two enzymes(41, 42). Therefore, we sought to determine what happens when a 1D-diffusing APE1 encounters pol ꞵ bound to a 1-nt gap. Here, we did not investigate pol ꞵ at AP-sites because of the low affinity of pol ꞵ for AP-sites compared with other BER DNA intermediates(41). We observed that in most cases (53%), motile APE1 bypassed the pol ꞵ-1nt gap complex without disrupting pol ꞵ (Fig. 6A-B). The next most common outcome (35%) was that diffusing APE1 could not scan past pol ꞵ, and the pol ꞵ-1nt gap complex remained intact. Only a minority of the time did APE1 evict pol ꞵ from the damage site (9%) or simultaneously bind the gap alongside pol ꞵ (2%). These results indicate that once pol ꞵ has bound to its DNA substrate, it is largely resistant to disruption by other BER proteins searching for damage. Notably, pol ꞵ can remain stably bound to the 1-nt gap even though we observe APE1 is capable of transiently binding the gap in the absence of a bound pol ꞵ (Supplementary Fig. 6A). At the same time, the presence of the pol ꞵ-1nt complex does not significantly impede APE1’s ongoing search along non-damaged DNA. These results suggest a mechanism where BER proteins can form stable complexes with DNA repair intermediates without interfering with each other’s search dynamics. This may be facilitated by motile BER proteins such as XRCC1 or APE1 conducting 1D-diffusion through hopping, allowing motile search that can more easily bypass bound proteins(39, 43).

**Figure 6.**
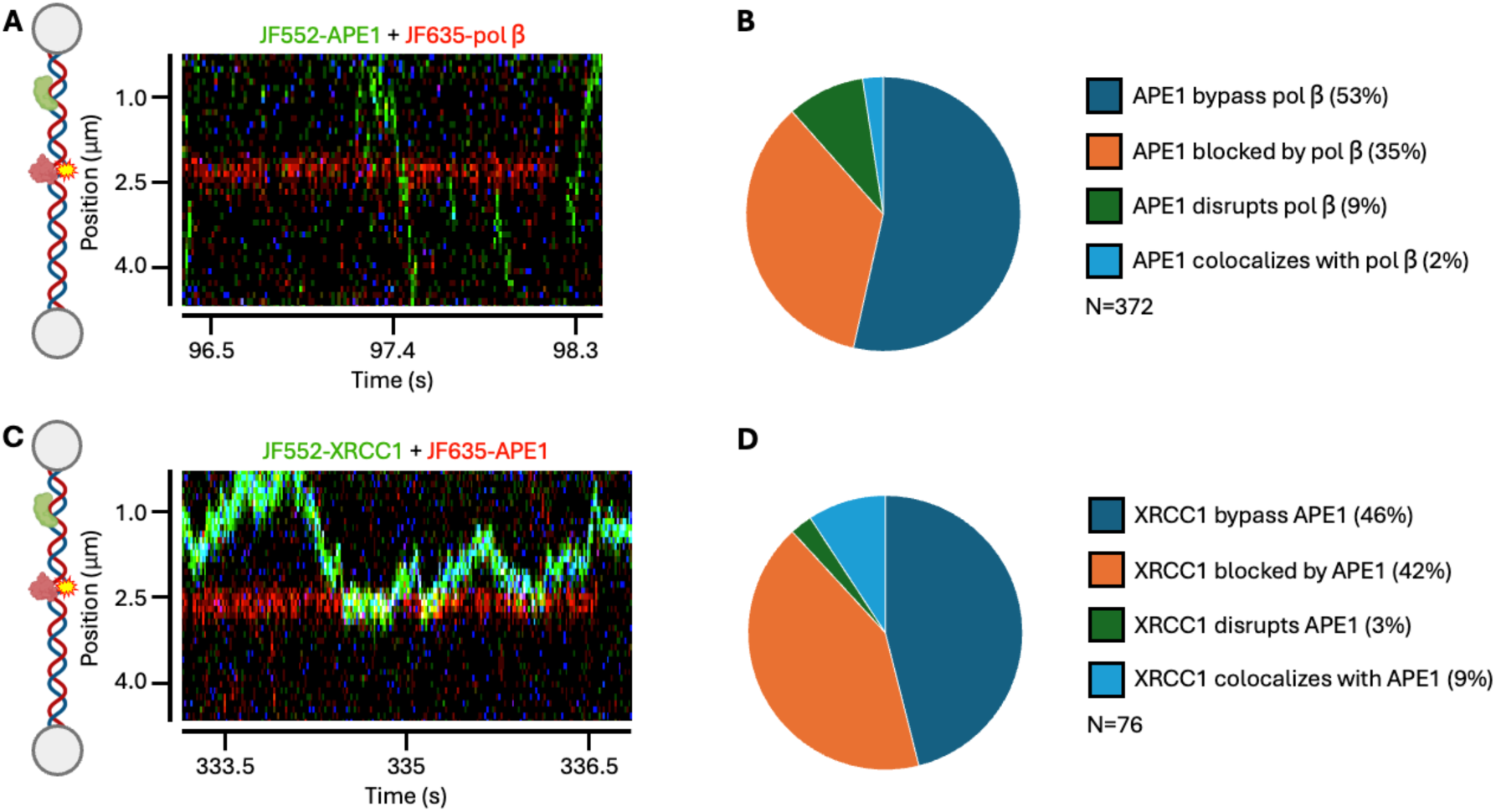
Motile BER proteins can continue DNA damage search past protein-bound DNA complexes. (A) Example kymograph of motile APE1 (green) searching through a pol ꞵ (red) bound to a one-nucleotide gap. (B) Percentage of motile APE1 events that were able to search through the pol ꞵ-1nt gap complex, were unable to search past the complex, disrupted the bound pol ꞵ, or colocalized with pol ꞵ at the 1-nt gap. Search through events required the mean intensity of the motile protein to pass ≥3 pixels beyond the stationary protein(C) Example kymograph of motile XRCC1 (green) unable to search through an APE1 (red) bound to an AP-site. (D) Percentage of motile XRCC1 events that were able to search through the APE1-AP complex, were unable to search through the complex, disrupted the bound APE1, or colocalized with APE1 at the AP-site.

We next asked whether the search mechanisms of APE1 itself may be disrupted by the presence of another motile BER protein, XRCC1. XRCC1 has previously been reported to interact with and modulate APE1 activity, though with lower affinity for APE1 than pol ꞵ(17, 44). To address this, we observed interactions between a non-motile APE1 bound to an AP-site and a motile XRCC1 undergoing 1D-diffusion along the DNA. Notably, we did not observe motile APE1-XRCC1 complex formation (in contrast to that observed with the pol ꞵ-XRCC1 complex). In most encounters, XRCC1 either bypassed the APE1-AP complex without perturbing either protein (46%) or failed to scan beyond APE1 (42%) (Fig. 6C-D). Like the prior case with a motile APE1 and stationary pol ꞵ, disruption of the protein bound at the damage site was uncommon. XRCC1 displaced APE1 from the AP site in only 3% of events and colocalized with APE1 at the lesion in 9% of cases. As previously reported, we observe XRCC1 was capable of binding the AP site in the absence of APE1, indicating that direct competition for this substrate is possible (Supplementary Fig. 6B)(15). Together, these results demonstrate that APE1 forms a stable complex at AP sites, that is largely resistant to displacement by other BER proteins engaged in damage search.

## Discussion

XRCC1 and pol ꞵ form a complex that is proposed to promote the recruitment of pol ꞵ to sites of DNA damage during base excision repair(21, 22, 45). However, unlike the well characterized and essential interaction between XRCC1 and LigIIIα, *in vivo* BER appears to proceed even when disrupting the interaction between XRCC1 and pol ꞵ, albeit at a lower efficiency(25). As a result, the precise functional role of XRCC1 as a scaffold protein during BER remains unclear. At the same time, pol ꞵ and other BER proteins must efficiently locate their specific DNA lesion while traversing through a huge excess of non-damaged DNA. How XRCC1 influences the process of pol ꞵ damage search, and whether its interaction with pol ꞵ contributes to lesion recognition or coordination with other repair steps is not known. Here, we used correlated optical tweezers with fluorescence microscopy to establish how pol ꞵ and XRCC1 search for DNA damage, the functional consequences of XRCC1 binding pol ꞵ during this search, and how motile BER proteins navigate DNA in the presence of other protein-DNA complexes.

Our results demonstrate that pol β binds both single-nucleotide gaps and 3′ nicks via non-motile (3D) diffusion. Notably, under our experimental conditions, we did not observe evidence of motile (1D) diffusion along non-damaged DNA preceding lesion engagement, similar to results reported elsewhere(28). We also observed the average dwell time at 1-nt gaps was approximately 3x longer than at the 3′ nick, consistent with pol ꞵ stable engagement and specificity for 1-nt gaps(41). Quantitative analysis of pol ꞵ at the 1-nt gaps revealed that increasing the tensile force on the DNA from 10 pN to 40 pN progressively reduced binding affinity through both a decrease in dwell time and increase in gap time. Consistent with this trend, the percentage of events in the long-lived binding population decreased with increasing force, with the affinity of pol ꞵ for the 1-nt gap approximately 14-fold lower at 40 pN than at 10 pN. In contrast, binding at the 3′ nick exhibited reduced dwell times and increased gap times from 10 pN to 20 pN, followed by a recovery at 40 pN. We speculate that moderate stretching at 20 pN may widen the 3′ nick into a structure that more closely resembled a 1-nt gap, thereby enhancing pol ꞵ binding(35), whereas further stretching distorts the lesion and reduces binding stability, possibly by mechanically opposing the ∼90° DNA bend induced by stable pol ꞵ binding(33). Together, these data support a model where pol ꞵ locates repair intermediates through direct 3D binding and where the mechanical conformation of the DNA plays a role in lesion recognition and binding stability.

We find that unlike pol ꞵ, XRCC1 readily binds non-damaged DNA and searches for lesions using motile diffusion. During an average binding event, XRCC1 was capable of scanning more than a quarter of the entire ∼12.6kbp DNA substrate. MSD analysis of XRCC1 traces further demonstrated that most motile search events were not rotationally-coupled, consistent with transient interactions with the phosphodiester backbone as XRCC1 scans for DNA damage (hopping). To our knowledge, this is the first direct demonstration that XRCC1 uses motile diffusion to locate DNA damage. In the presence of either a single-nucleotide gap or 3′ nick, XRCC1 can bind directly to the damage site via 3D-diffusion or locate the lesion by first binding non-damaged DNA and subsequently searching via 1D-diffusion. Notably, in the presence of a 1-nt gap, XRCC1 bound the damage only 14% of the time, with 80% of all events undergoing motile diffusion, similar to the behavior of XRCC1 in the absence of damage. These observations suggest that XRCC1 may function primarily to facilitate efficient lesion search rather than prolonged retention at DNA damage. Specifically, the ability to access both 1D and 3D modes of diffusion is likely to enhance the efficacy of XRCC1 damage search. In this model, the 1D-diffusion allows XRCC1 to interrogate local DNA without dissociating, essentially confining the protein to a region of the genome. In contrast, 3D-diffusion enables XRCC1 to sample more distant sites and to bypass protein-DNA obstacles that would otherwise limit continuous 1D scanning.

When pol ꞵ and XRCC1 form a stable complex, they exhibit predominantly motile (1D) diffusion on non-damaged DNA, enabling pol ꞵ to scan DNA through a mechanism not observed alone. The dwell time of the complex on non-damaged DNA closely matched that of XRCC1 alone, suggesting that DNA engagement and search behavior of the complex are primarily driven by XRCC1(46). Consistent with this, the complex displayed largely non-rotationally coupled 1D-diffusion, with a median diffusivity roughly 2.5-fold lower than that observed for XRCC1 alone. In the presence of DNA damage, complex formation allowed pol ꞵ to access both motile and non-motile diffusion to locate 1-nt gaps and 3′ nicks. Notably, the dwell times of the complex at gaps were similar to those of pol ꞵ alone, indicating that lesion specificity within the complex is dominated by pol ꞵ rather than XRCC1. Furthermore, in the majority of cases pol ꞵ and XRCC1 bound to and dissociated from damage sites as a pre-formed complex. However, when the complex assembled or disassembled at the lesion, pol ꞵ more frequently bound first, whereas XRCC1 was more likely to dissociate first. This pattern supports a model where XRCC1 facilitates the search for DNA damage but is not primarily responsible for enhancing pol β binding at the lesion, consistent with prior biochemical studies(20, 22). Instead, XRCC1 may function to locate damage and retain pol β in the general region where damage is present. Additionally, because XRCC1 functions as a scaffold for LigIIIα recruitment, these transient XRCC1 binding events at damage may reflect attempts to coordinate downstream repair following gap filling. More broadly, this represents one of the first examples of a protein complex altering the search behavior of a DNA-binding enzyme by promoting motile diffusion on DNA(47).

We further demonstrated that motile protein search in BER rarely disrupted pre-existing protein-DNA complexes. Motile APE1 was unlikely to displace pol ꞵ bound at a single-nucleotide gap, even though these proteins are known to interact(41). This indicates that pol ꞵ can stably occupy the lesion while other BER proteins search past it, disrupting neither the damage bound complex nor the motile protein. A similar observation was made with XRCC1, where the motile XRCC1 infrequently disrupted APE1 bound to an AP-site, suggesting that these complexes are likewise stable in the presence of searching proteins. Together, these results indicate that BER proteins can search through genomic regions where repair is actively occurring without destabilizing existing protein-DNA complexes, consistent with a hopping mechanism of diffusion allowing motile proteins to bypass these complexes more easily than by continuous sliding along the helix(48). Such a mechanism would enable BER factors to navigate crowded chromatin environments while preserving pre-bound intermediates. In this model, searching proteins can accumulate within the local region of repair but continue to search through each other, promoting efficient substrate handoff and enabling multiple steps of BER to occur within a shared local DNA environment(42).

Taken together, our findings support a model in which DNA damage search by XRCC1 and pol ꞵ is altered by complex formation, resulting in a more adaptive damage search that combines complementary diffusion strategies of each protein (Fig. 7). Pol ꞵ primarily samples DNA damage through 3D diffusion from bulk solution, enabling detection of distantly spaced lesions, but which is inefficient in searching locally within a defined genomic region. XRCC1 can also locate damage through non-motile diffusion, but is capable of 1D diffusive motile search, allowing it to scan nearby DNA. Formation of the pol ꞵ-XRCC1 complex integrates both search strategies, enabling pol ꞵ to access both local and long-range damage search mechanisms. In this model, motile 1D scanning is driven largely by XRCC1, whereas lesion recognition and retention are dominated by pol ꞵ. This suggests that XRCC1 functions primarily to extend the local damage search of pol ꞵ rather than to stabilize the pol ꞵ-DNA complex once a lesion has been identified. This may allow XRCC1 to retain pol ꞵ in areas of oxidative damage where BER is occurring, but where a pol ꞵ substrate is not yet available. Such pre-positioning of pol ꞵ on non-damaged DNA near repair sites while upstream BER steps including base excision and AP site incision are being completed, would enhance the ability of pol ꞵ to find the gap and enhance overall repair efficiency(49).

**Figure 7.**
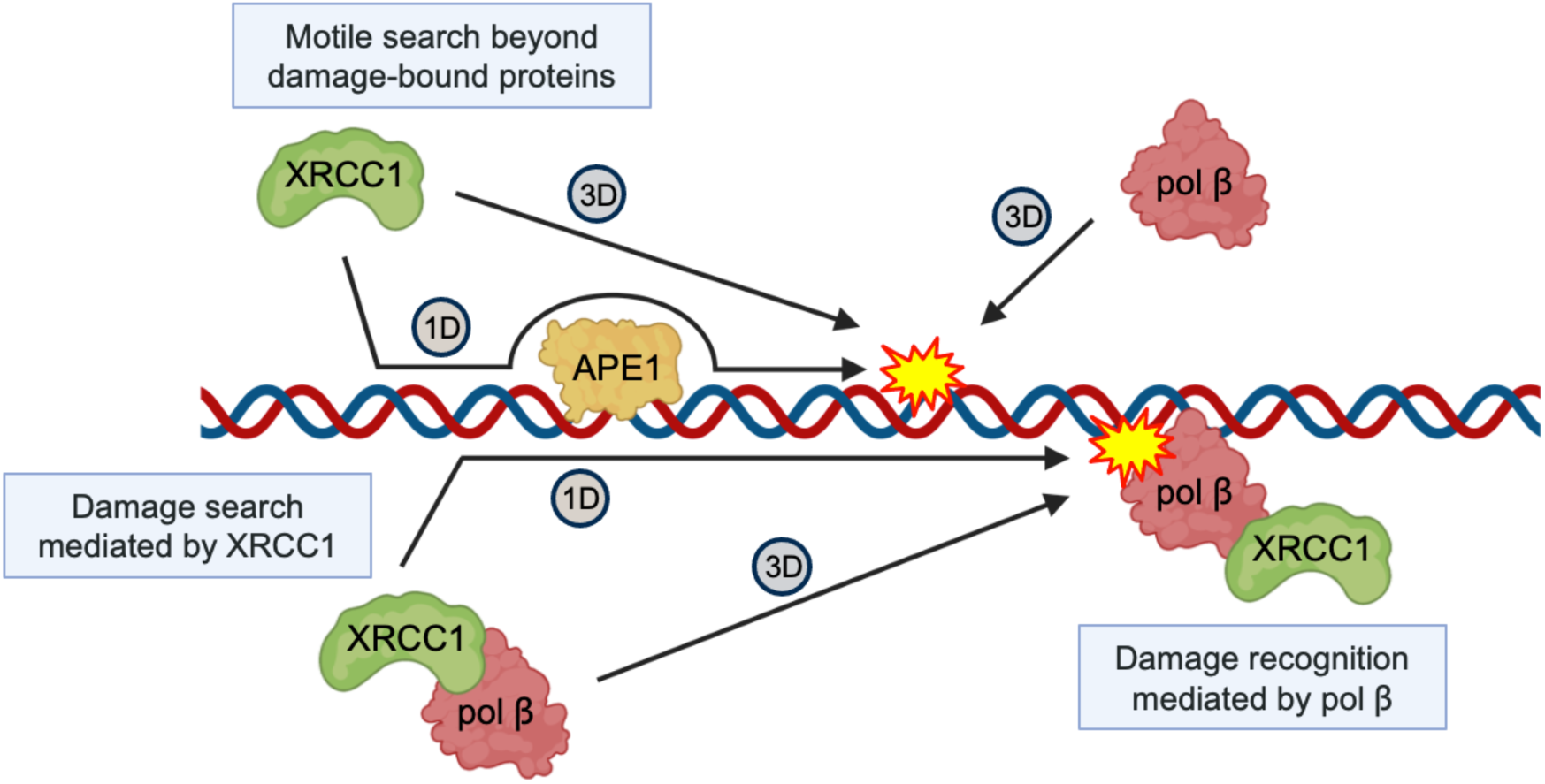
Model of pol ꞵ, XRCC1, and the pol ꞵ/XRCC1 complex locating DNA damage. We propose that pol ꞵ locates damage using 3D-diffusion without extensive binding at non-damaged DNA. XRCC1 can also undergo 3D-diffusion to locate damage but can also engage non-damaged DNA to search via 1D-diffusion and scan past other protein-DNA complexes during search. Formation of the pol ꞵ-XRCC1 complex allows both pol ꞵ and XRCC1 to use either mode of diffusion to locate damage. Complex search by 1D-diffusion primarily relies on XRCC1 while pol ꞵ is primarily responsible for recognition of the damage site. Made with Biorender.com

It is also interesting to consider these results in the context of PARP1 and auto-PARylation, which facilitates the recruitment of XRCC1 and the subsequent assembly of downstream BER factors including pol β and LigIIIα(50, 51). Prior work has proposed a role for XRCC1 in limiting excessive PARP1 engagement at sites of damage and promoting the assembly of downstream repair complexes, thereby facilitating PARP1 release following PARylation(12, 52). Recent cellular work has extended this model, demonstrating that the essential, non-redundant role of both pol β and XRCC1 during BER is to suppress excessive PARP1 engagement at repair intermediates, a function the two proteins perform together and that requires their direct interaction(24). The 1D-diffusion capacity of XRCC1 observed here offers a plausible mechanism for this coordination: upon PARP1-mediated recruitment to the damage site, XRCC1 may actively scan local chromatin to identify repair intermediates and coordinate downstream BER factor assembly, including pol ꞵ, in a manner that facilitates timely PARP1 release and prevents cytotoxic PARP1 trapping. The accelerated delivery of pol β to incised intermediates by the motile complex would narrow the window for PARP1 over-engagement, providing a mechanistic basis for this cellular requirement. Consistent with a shared interface underlying both phenomena, the XRCC1-F67A mutant that cannot bind pol β fails to fully support rapid cellular BER(24), while the corresponding pol β triple mutant (L301R/V303R/V306R) abolished formation of the motile complex in our experiments (Supplementary Fig. S5). Together, these findings place the search behaviors of pol ꞵ and XRCC1 in a broader context of BER coordination, where the interplay between motile and non-motile diffusion mechanisms, scaffold-driven retention, and simultaneous damage search defines the organization and efficiency of repair complexes at DNA damage.

## Materials and Methods

### Halo Protein Expression, Purification, and Fluorescent Labeling

We expressed N-terminally Halo-tagged pol ꞵ in BL21 *E. coli* using the pFN29A vector (Promega). C-terminally Halo-tagged XRCC1 and APE1 were expressed using the pFC30K vector (Promega). Bacteria were grown using 2XYT media at 37°C in a bioreactor until reaching an optical density at 600nm of 0.6. Protein expression was induced by addition of IPTG to the media and cultures allowed to grow at 18°C for 18 hours. Cells were harvested and lysed by sonication using a Fisher Scientific FB120 sonicator for 12 cycles of 50 seconds at 90% power. Lysates were clarified from solid products by ultracentrifugation at 24,000xg for 1 hour. Halo-tagged pol ꞵ was purified by loading clarified lysate onto an ÄKTA Pure FPLC system with a Cytiva 5mL His affinity column. Resulting fractions containing the Halo-pol ꞵ were combined and further run on a Resource Q anion exchange column to remove contaminating DNA. These fractions were then loaded onto a Cytiva Superdex 200 Increase 10/300 gel filtration column. A similar purification strategy was used to isolate XRCC1-Halo and APE1-Halo. Subsequently, purified Halo-pol ꞵ, XRCC1-Halo, or APE1-Halo were fluorescently labeled by incubating 10 nmol of protein with 20 nmol of Janelia Fluor 552 or Janelia Fluor 635 Halo-Tag dyes for 1 hour at room temperature(53). Labeled proteins were further purified using an ÄKTA-Pure FPLC with a Cytiva Superdex 200 Increase 10/300 gel filtration column. Labeling efficiency was determined by UV-vis spectroscopy to measure the ratio of protein concentration to dye concentration in our purified fractions. Labeling efficiency for all Halo-pol ꞵ, XRCC1-Halo, and APE1-Halo was >80% for all samples. Sample aliquots were then stored at −80°C until use.

### DNA Substrate Ligation and Tethering

DNA substrates for CTFM experiments were generated by ligating pre-annealed DNA oligonucleotides (Integrated DNA Technologies) between two ∼6.3 kbp DNA handles generated by Lumicks. Each DNA handle is biotinylated at one end and contained either an Atto647N fluorophore with a GTTG single-strand overhang or an Atto488 fluorophore with a TGGT overhang. A ∼50bp DNA insert containing either an internal 1-nt gap, 3*′*-OH nick, or a non-hydrolyzable AP-site was annealed and ligated between the two DNA handles(54). Ligation reactions were performed according to the Lumicks DNA tethering protocol. Briefly, 0.05 pmol of annealed insert oligos were combined with the DNA handles, DNA ligase buffer, and T4 DNA ligase. These reactions were incubated at room temperature for 16 hours and subsequently stored at 4°C. For substrates containing the 3′-OH nick, T4 DNA ligase was heat inactivated at 70°C for 10 min. These were then incubated with 1mM ATP, PNK reaction buffer, and T4 PNK to phosphorylate the 5′-OH of the nicked substrate prior to CTFM experiments.

### Correlated Optical Tweezers with Fluorescence Microscopy (CTFM)

All CTFM experiments were performed using a Lumicks C-Trap instrument equipped with a microfluidics system, dual optical traps, and a three-color confocal fluorescence microscope with excitation lasers at 488, 561, and 638 nm. Prior to each experiment, the microfluidics system and flow cell were prepared according to Lumicks protocols. Briefly, approximately 0.5 mL of 0.1% bovine serum albumin (BSA) was flowed through each channel and incubated for 30 min to passivate the system. This was followed by incubation with approximately 0.5 mL of a 1:10 diluted Pluronic solution for an additional 30 minutes. Halo-tagged protein samples were prepared by serial dilution of previously frozen aliquots into 0.45 µm filtered C-Trap buffer (20 mM HEPES, pH 7.2, 100 mM NaCl, 1 mM EDTA, 1 mM TCEP, 0.1 mg/mL BSA, and 1mM Trolox). Lumicks streptavidin-coated 1.5-1.9 µm polystyrene beads diluted to 1:1333 in C-Trap buffer were loaded into channel 1. DNA substrates diluted 1:250 in C-Trap buffer were loaded into channel 2, C-Trap buffer alone into channel 3, diluted Halo-tagged protein into channel 4, and C-Trap buffer only into channel 5. Following sample loading, two polystyrene beads were captured in channel 1 using the optical trapping lasers and used to generate bead templates. Fresh beads were then captured and moved to channel 2, where the beads bound a single biotinylated DNA. To verify single DNA tether formation, bead pairs were transferred to channel 3 which contained only buffer. Using the optical traps, one bead was displaced while the other was kept stationary to apply tensile force along the DNA tether. A force-distance curve was generated by this action, and compared to a worm-like chain (WLC) model corresponding to the stretching of a ∼12.6 kbp double stranded DNA(55). After verifying only one DNA was present, fluid flow was stopped, the DNA tether stretched to 10-40 pN based on the WLC model, and the DNA tether was transferred to channel 4 containing fluorescently labeled protein(s). Confocal kymographs were acquired with a pixel size of 0.1 µm, a line time of 15 ms, and excitation laser powers set to 5%. All experiments were conducted at room temperature with multiple kymographs collected on separate days from independent DNA ligation preparations.

### Kymograph Analysis and Particle Tracking

Kymographs were collected using Lumicks Bluelake software and exported as .h5 files. Subsequent analyses were performed in Python 3.12.2 with the Lumicks PyLake package version 1.5.2. Anaconda 24.7.1 was used for environment management and Jupyter Notebook 7.0.8 served as the integrated development environment. Particle tracking was performed using the Kymotracking widget features in Pylake, which implements the Lumicks KymoGreedy algorithm to identify the tracking lines. Customizable algorithm parameters were adjusted to assign the best track to each individual trace collected.

### Calculating Particle Dwell Time and Gap Time

Dwell times for a single protein binding event binding the DNA substrate were defined as:

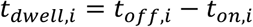

Where *t_on_*_,*i*_ is the time at which the *i*-th binding event begins and *t_off_*_,*i*_ is the time at which the same event ends. Similarly, the gap time between protein binding events was defined as the interval between the end of one binding event and the beginning of the next binding event:

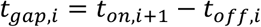

where *t_on_*_,*i*+1_ is the start of time of the subsequent protein binding event. To characterize distributions of dwell and gap times, cumulative distribution function (CDF) plots were constructed, defined as the survival probability *S*(*t*) = *P*(*T* ≥ *t*). These distributions were fit to single-, double-, and triple-exponential decay functions of the general form:

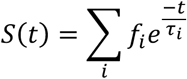

where *f_i_* represents the fractional contribution of each kinetic population and *τ_i_* is the corresponding lifetime. The sum of all fractional contributions was constrained such that ∑*_i_ f_i_* = 1. From these fits, a weighted average lifetime was calculated as:

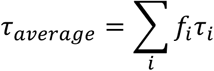

An effective off-rate and on-rate were then calculated from the average weighted binding lifetime and gap interval, respectively:

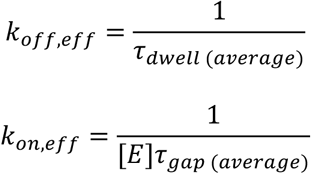

where [E] represents the enzyme concentration. From these, an effective binding affinity was determined from the ratio of the effective rate constants:

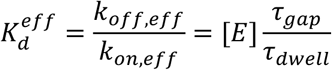

### Particle Diffusivity and Mean Squared Displacement

Mean squared displacements (MSD) was calculated for each track exhibiting motile behavior using the following equation:

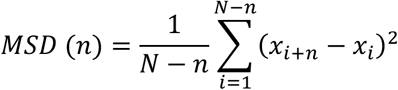

where *N* is the number of frames in the trajectory, *n* is the number of frames at a given step time, and *x_i_* is the particle position in the *i*-th frame(28, 56). Subsequently, the diffusion coefficient (*D*) was determined by fitting the linear portion of the MSD versus time plots to a one-dimensional diffusion model:

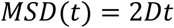

Fits were manually adjusted to include the linear portion of the graphs with fits resulting in an R^2^ < 0.8 or using <10% of the MSD plot were excluded.

The theoretical maximum diffusion coefficient for one-dimensional diffusion without rotationally-coupled motion was estimated using the Stokes-Einstein relation(57):

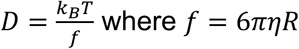

where *k_B_* is the Boltzmann constant, T is the system temperature, *η* is the viscosity of the aqueous medium, estimated as 9 x 10^-31^ Js/nm^3^, and *R* is the hydrodynamic radius of the Halo-protein or Halo-protein complex. *R* was estimated as 6.11 nm for XRCC1-Halo and 7.16 nm for the Halo-pol ꞵ/XRCC1-Halo complex using HullRad and structures of both proteins generated using AlphaFold v3(58, 59). We also calculated the theoretical maximum limit of one-dimensional diffusion for rotationally-coupled diffusion using the following equation:

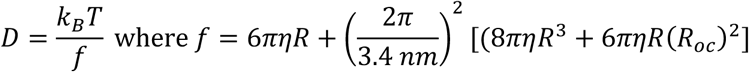

where *R_oc_* is the sum of the hydrodynamic radius of the protein or protein complex and the distance of the protein to the DNA center. *R_oc_* was estimated as 7.11 nm for XRCC1-Halo and 8.16 nm for the Halo-pol ꞵ/XRCC1-Halo complex. The helical pitch of DNA was taken as 3.4 nm per turn.

The median scan length was estimated from the root mean squared displacement:

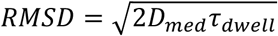

where *D_med_* is the median diffusion coefficient and *τ_d_*_w_*_ell_* is the average weighted binding lifetime as defined above.

## Acknowledgments

We thank Tyler M. Weaver and Bennett Van Houten for their contributions to project conceptualization and methodology and for critical reading of the manuscript. This work was supported by National Institutes of Health grants R35GM128562 to B.D.F. and F31ES036459 to S.H.T. The content is solely the responsibility of the authors and does not necessarily represent the official views of the National Institutes of Health.

## Data Availability

Further information and requests for reagents should be directed to the corresponding author.

## Author Contributions

Spencer H. Thompson was responsible for project conceptualization, methodology, investigation, data curation, data analysis, data presentation, original manuscript drafting, and manuscript editing. Kaitlin M. DeHart was responsible for investigation, data analysis, and manuscript review and editing. Matthew A. Schaich was responsible for methodology, data analysis, and manuscript review and editing. Bret D. Freudenthal was responsible for project supervision, project conceptualization, manuscript review and editing.

