## Supplemental Information for "XRCC1 Enables the Efficient Local Search for DNA Damage by DNA Polymerase Beta"

**A**

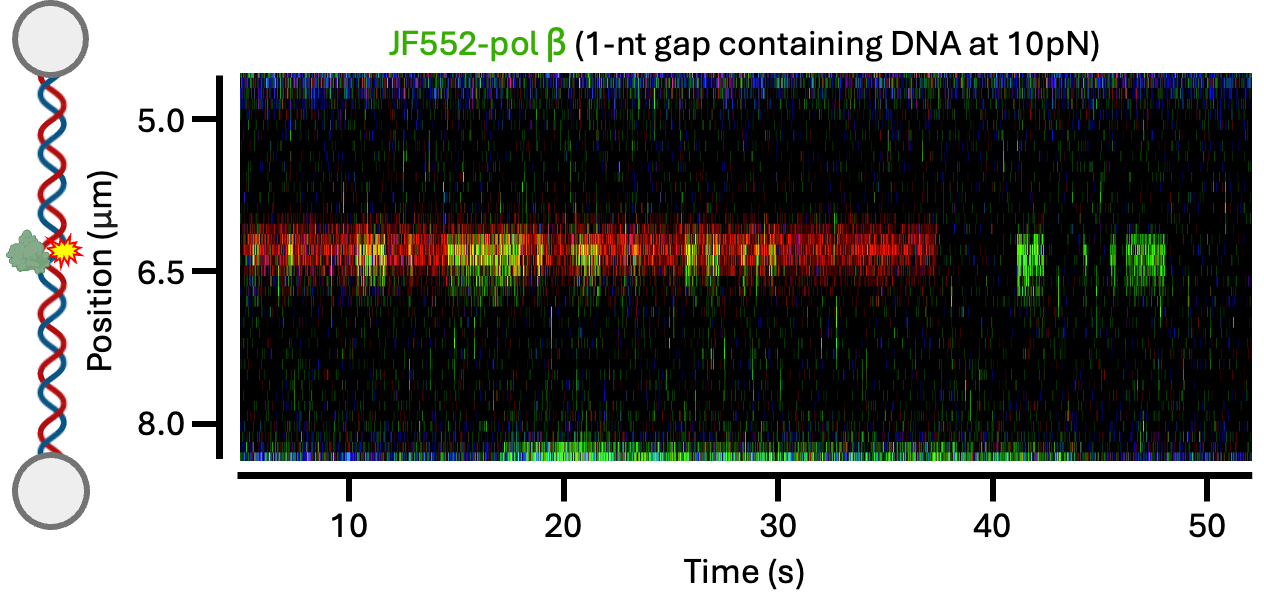

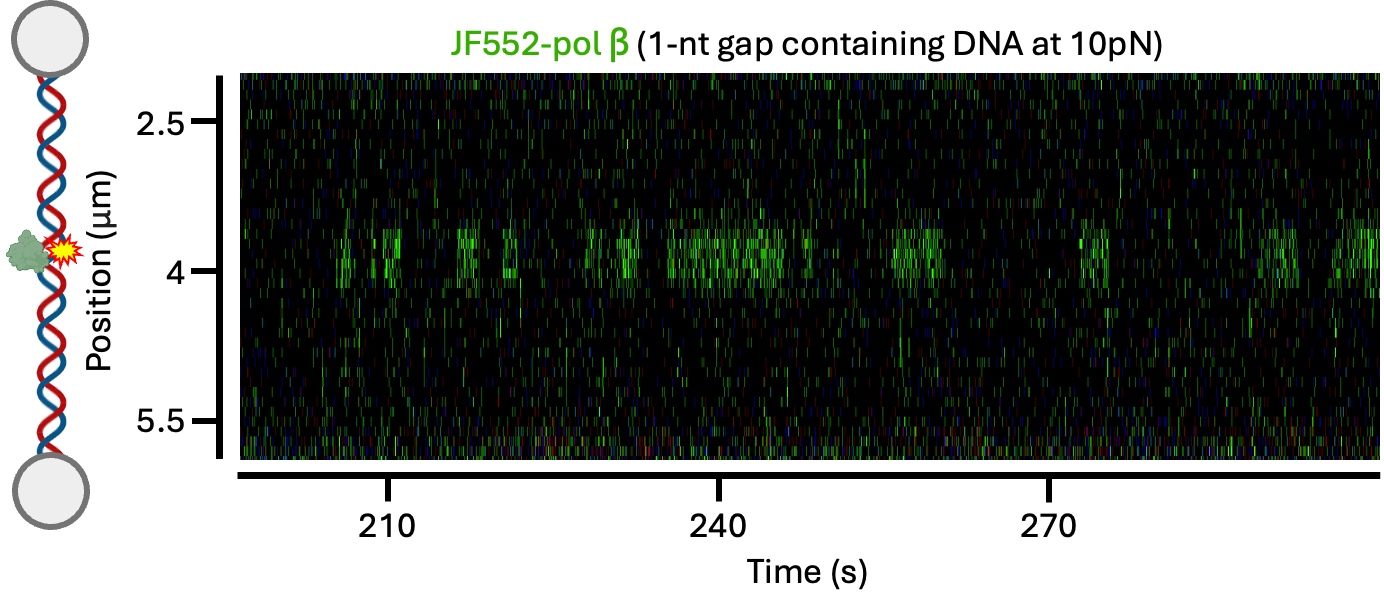

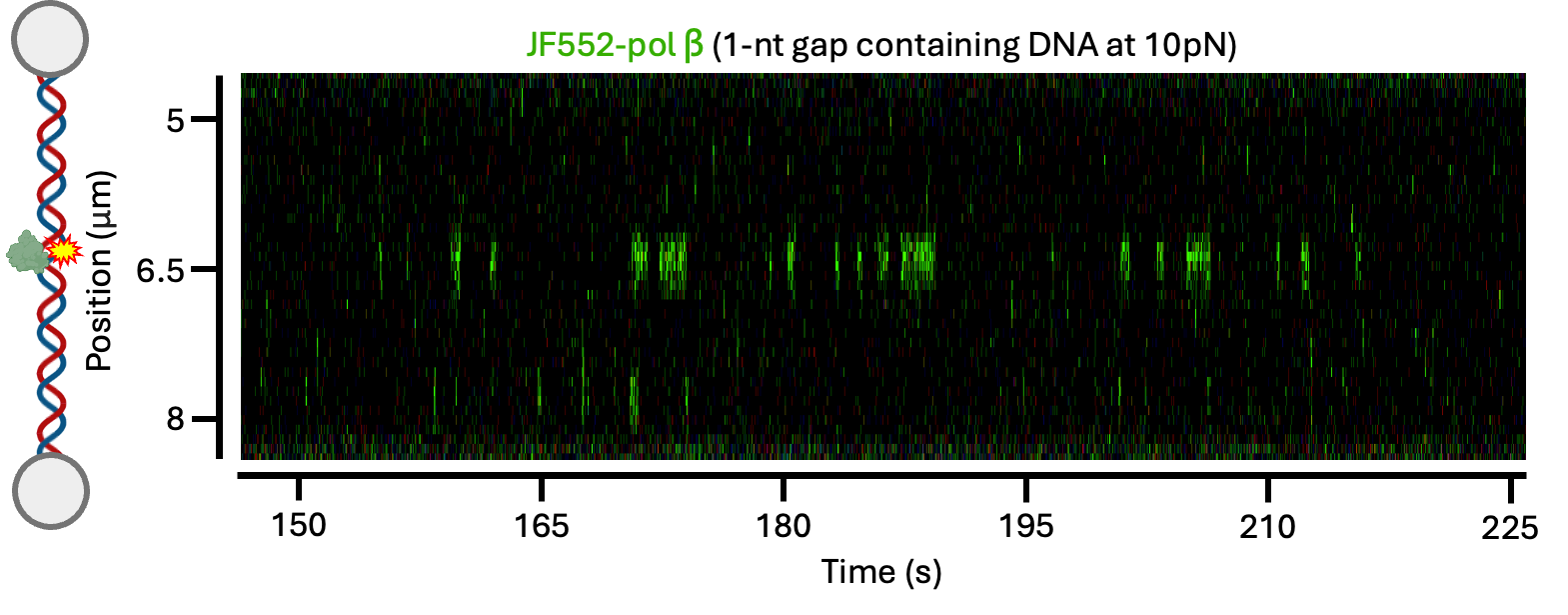

**B**

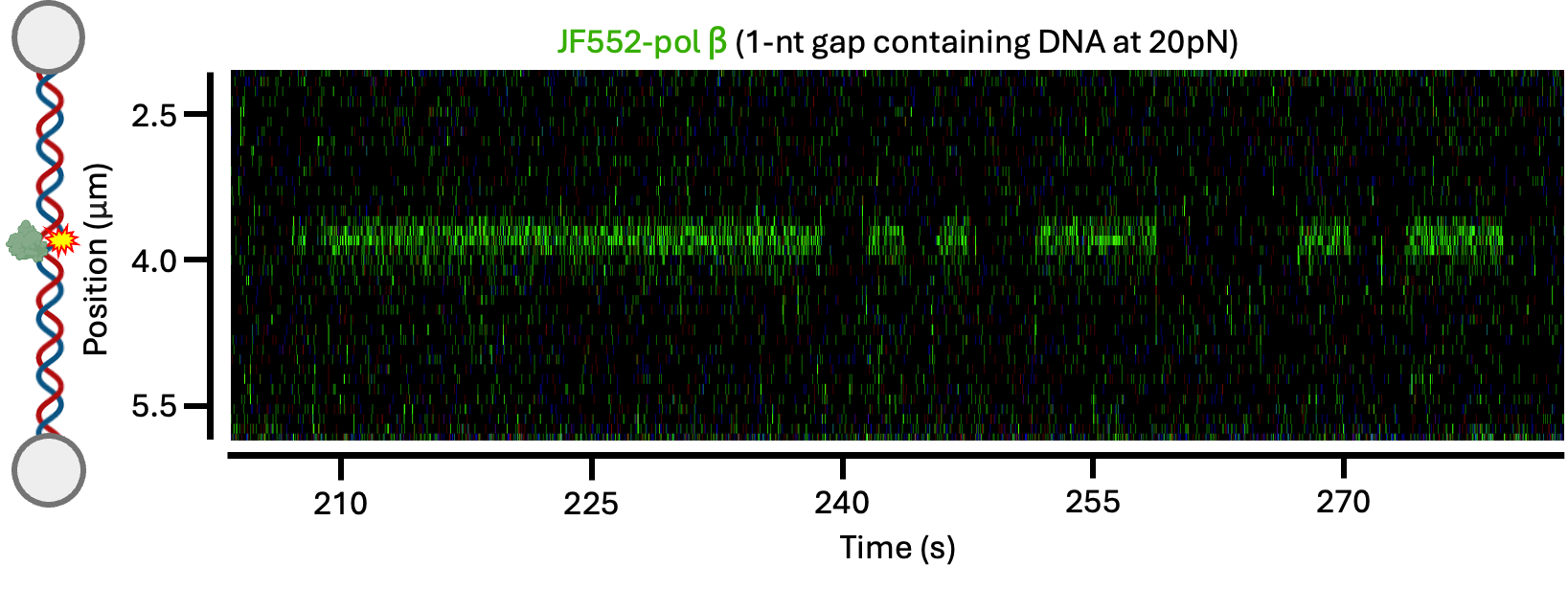

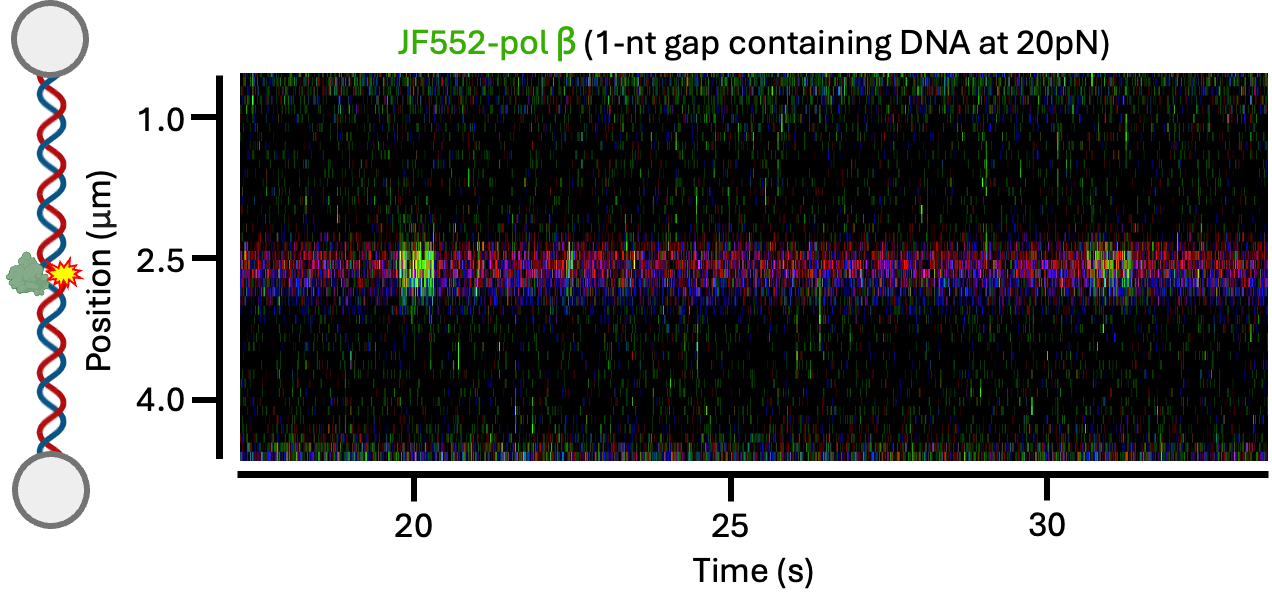

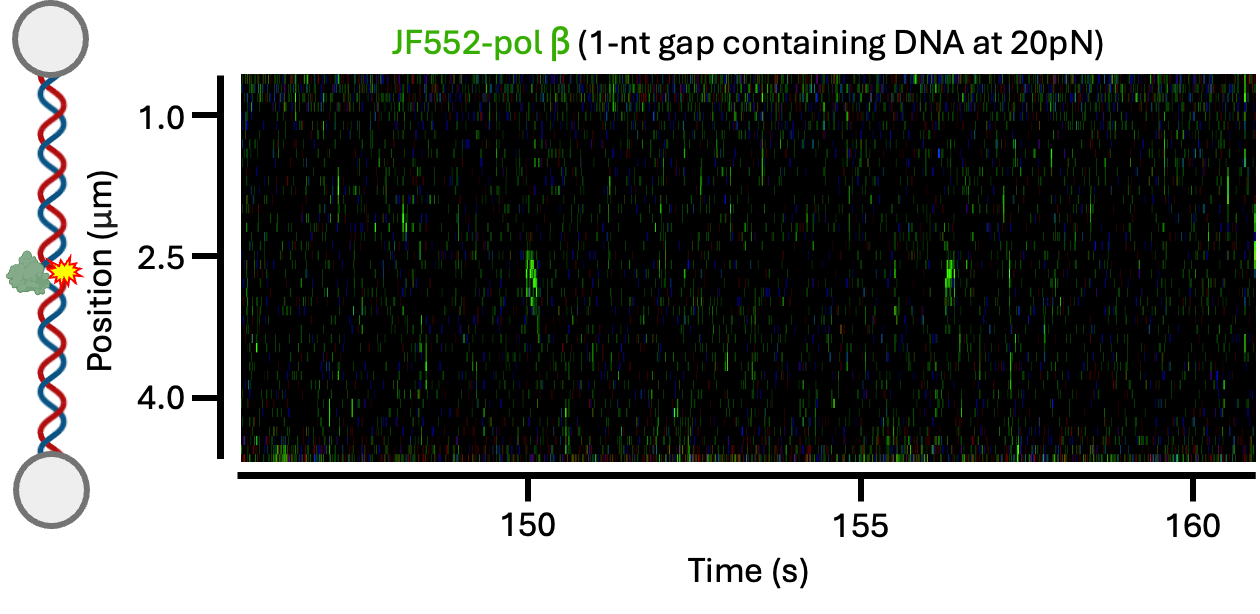

**C**

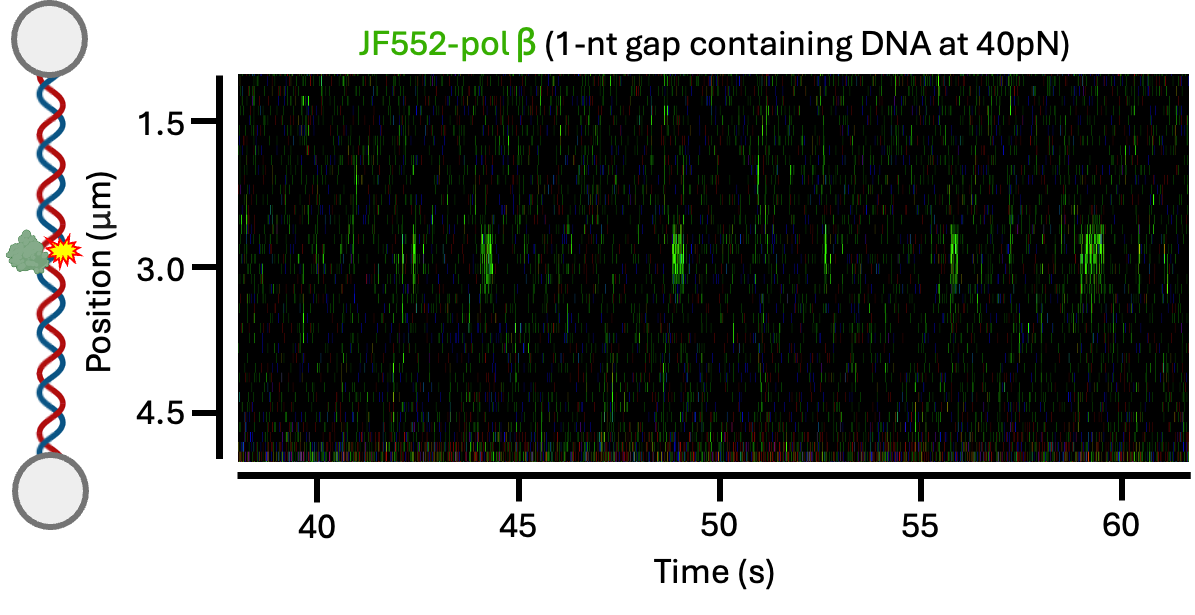

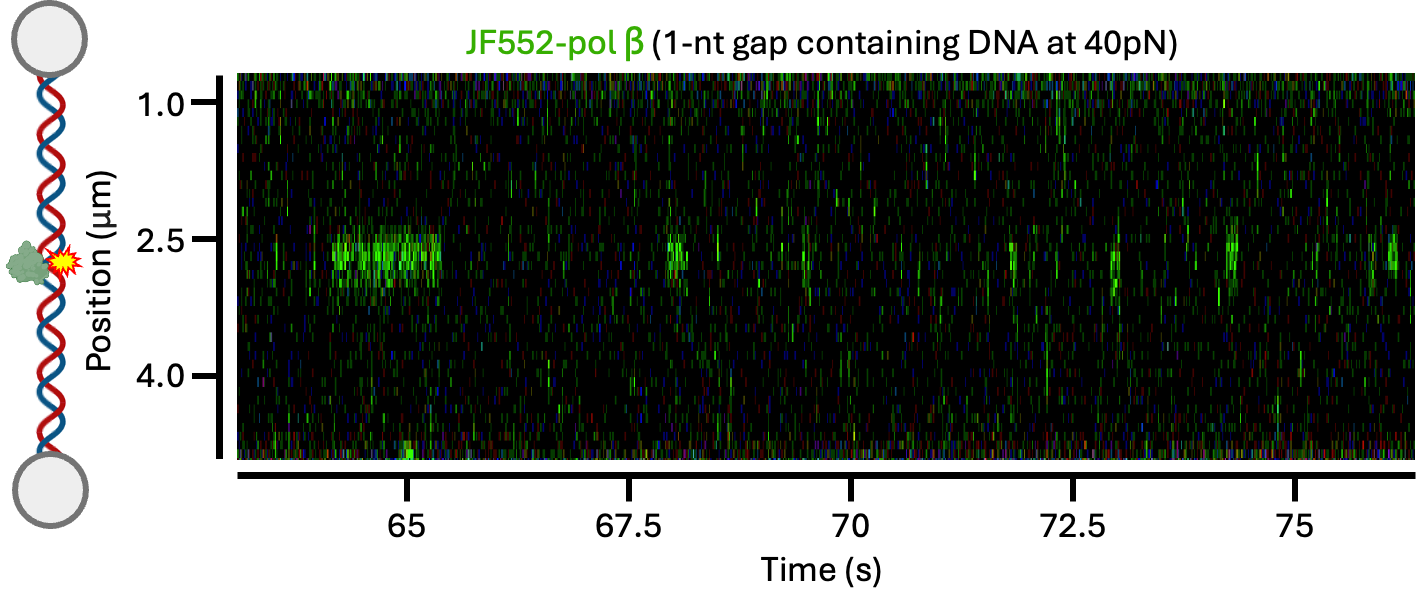

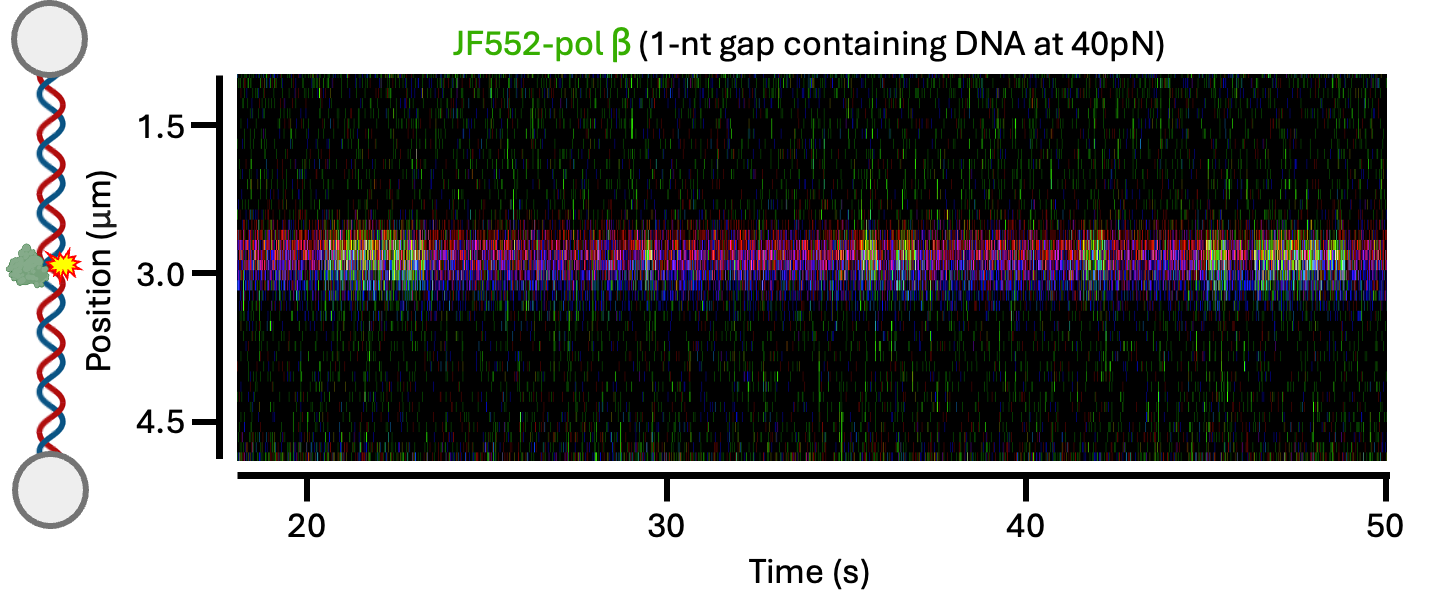

**Supplementary Figure S1. Additional kymographs of pol ꞵ on 1-nt gapped DNA at different forces**. (A) Example kymographs showing JF552-pol ꞵ binding to the gap stretched to 10 pN (B) JF552-pol ꞵ binding to the gap stretched to 20 pN. (C) JF552-pol ꞵ binding to the gap stretched to 40 pN.

**A**

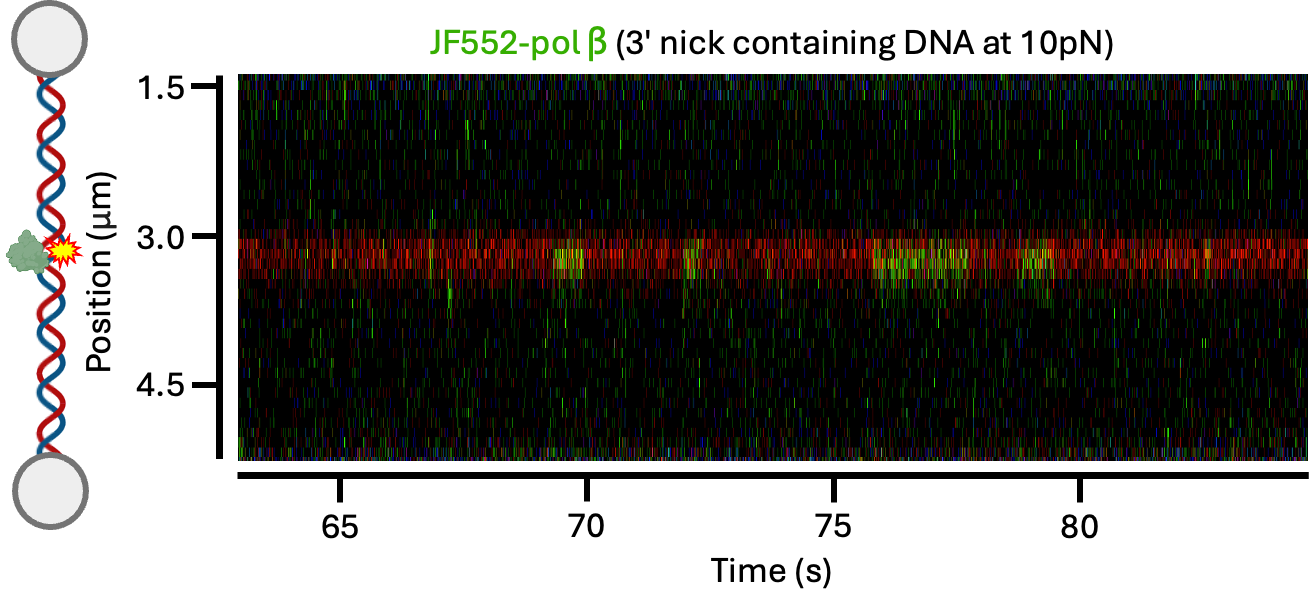

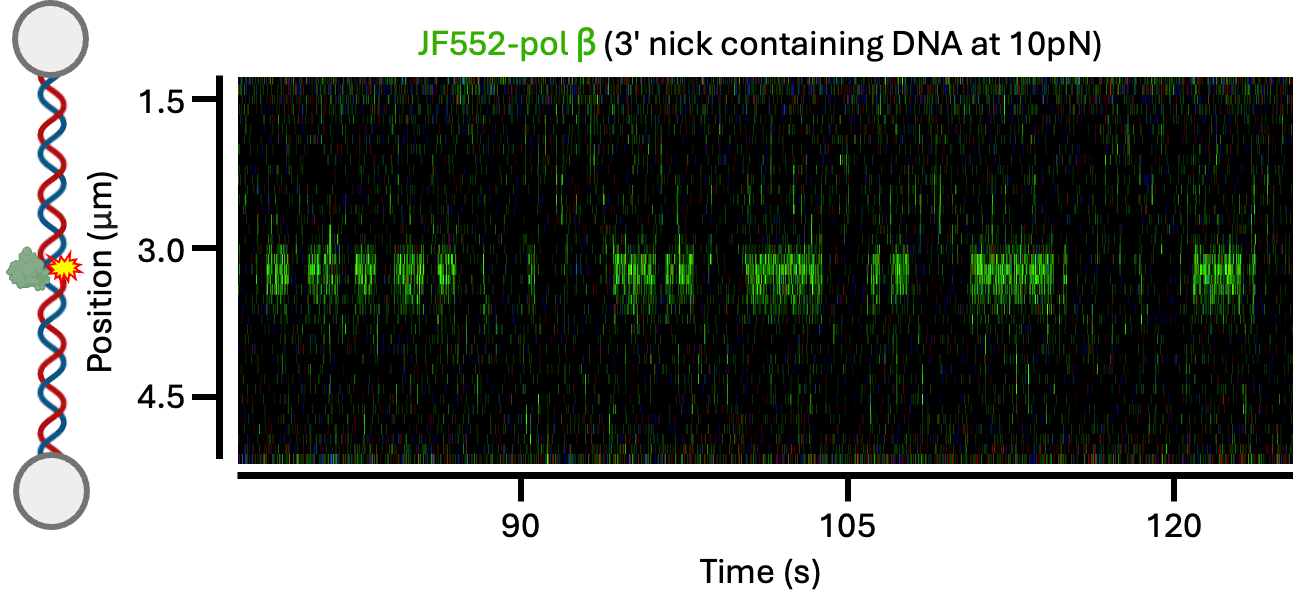

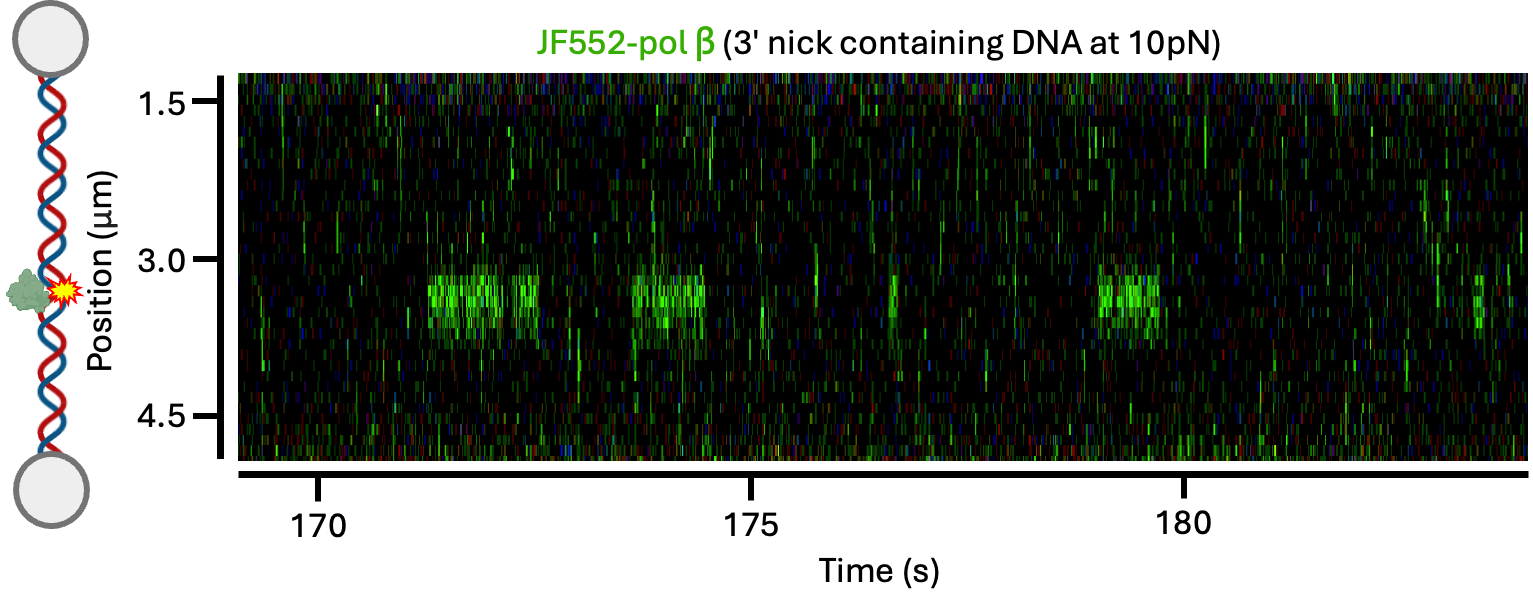

**B**

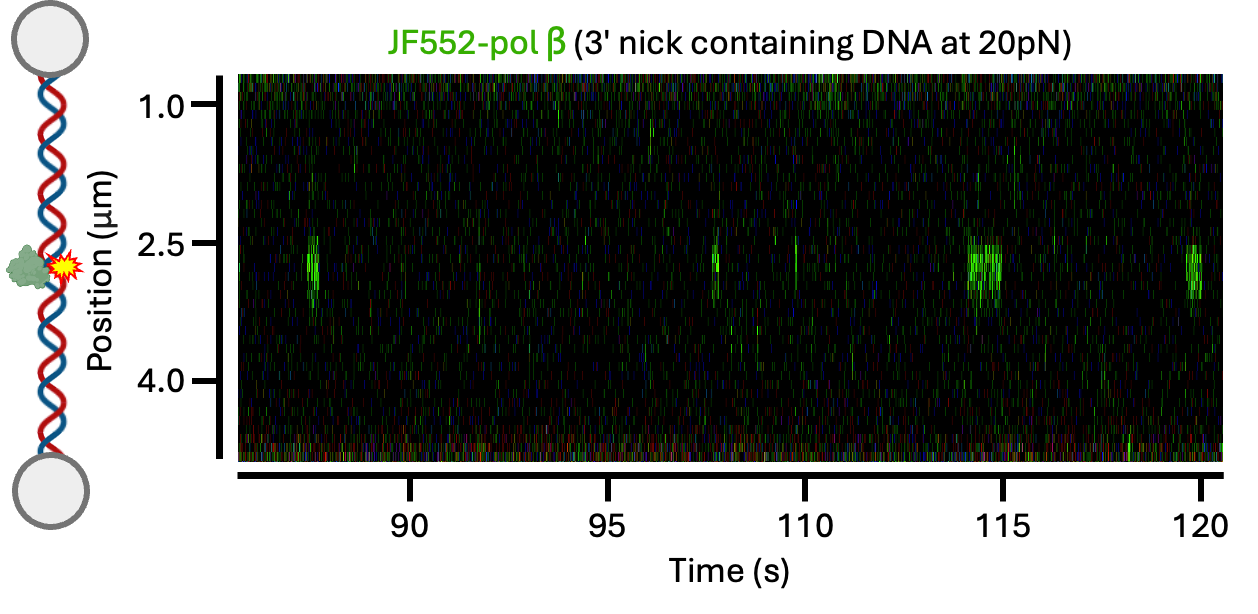

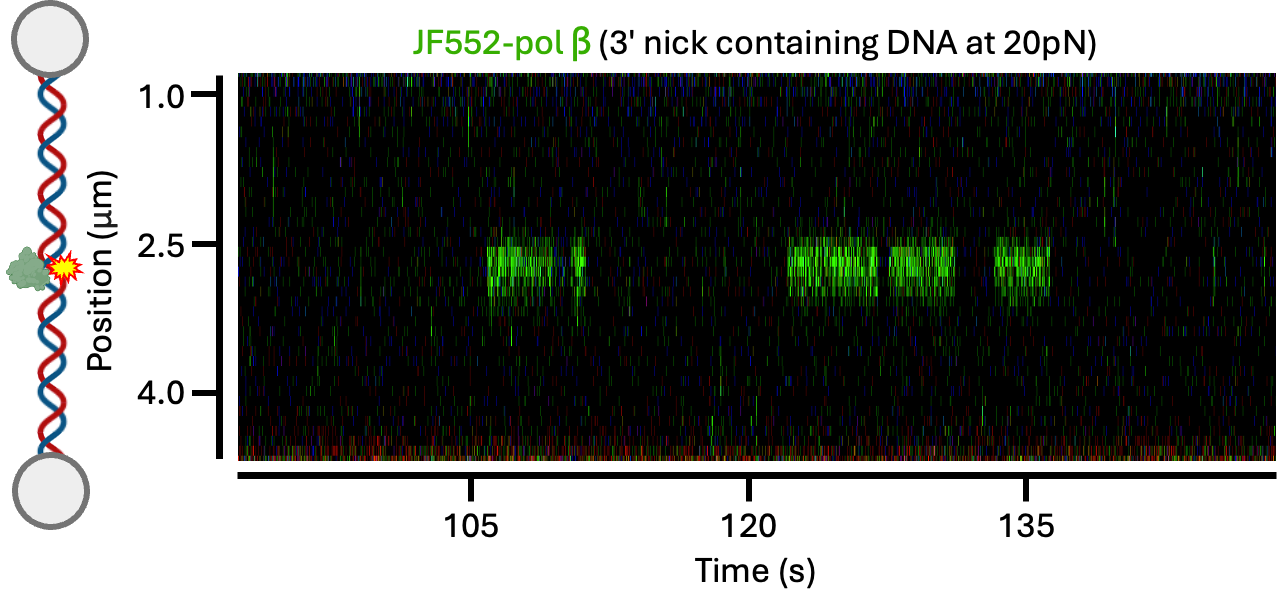

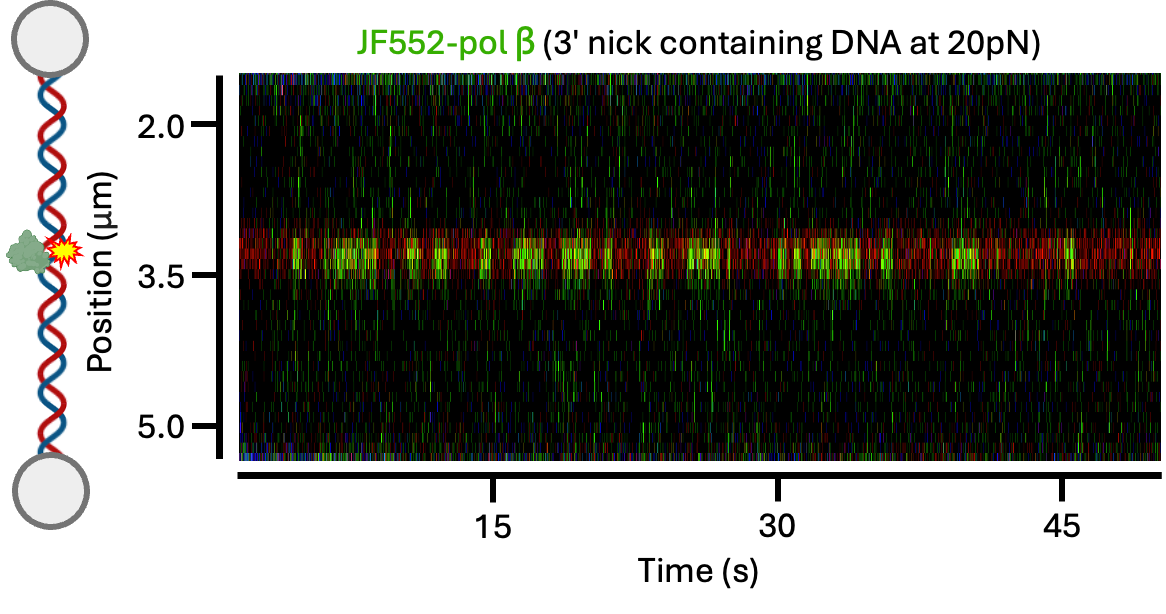

**C**

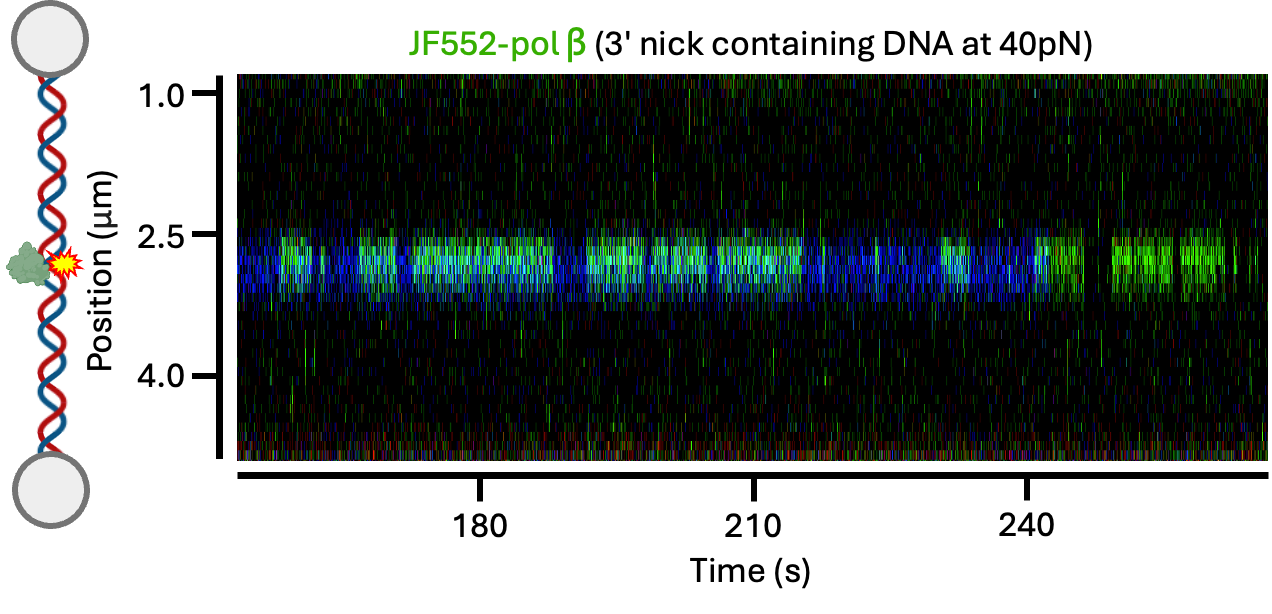

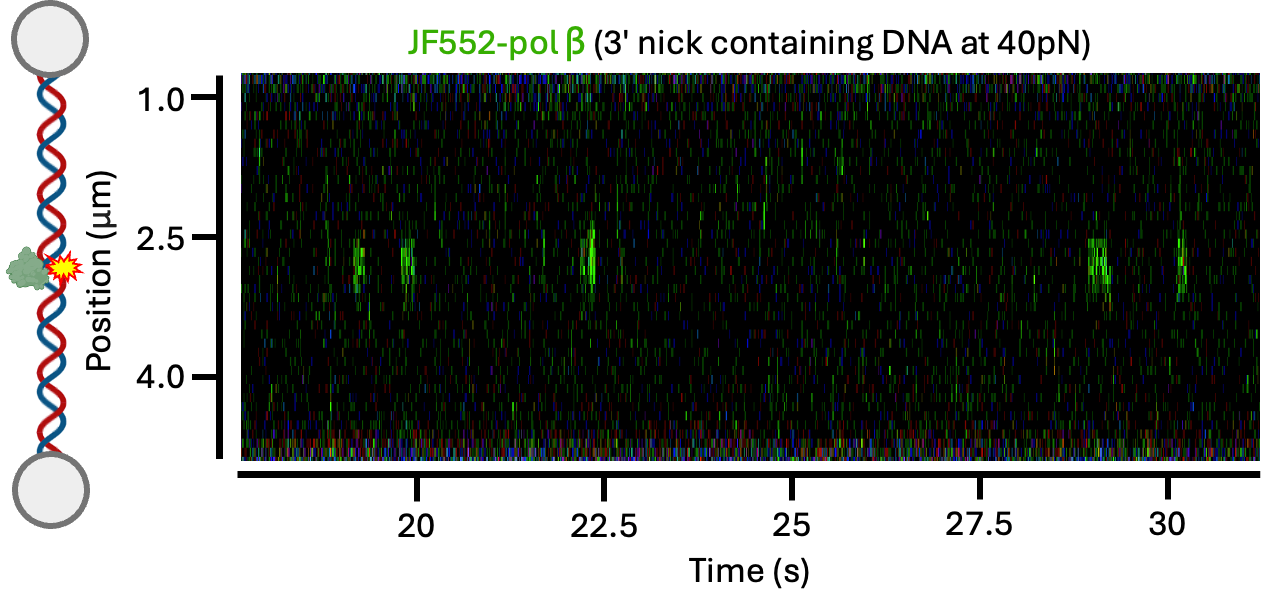

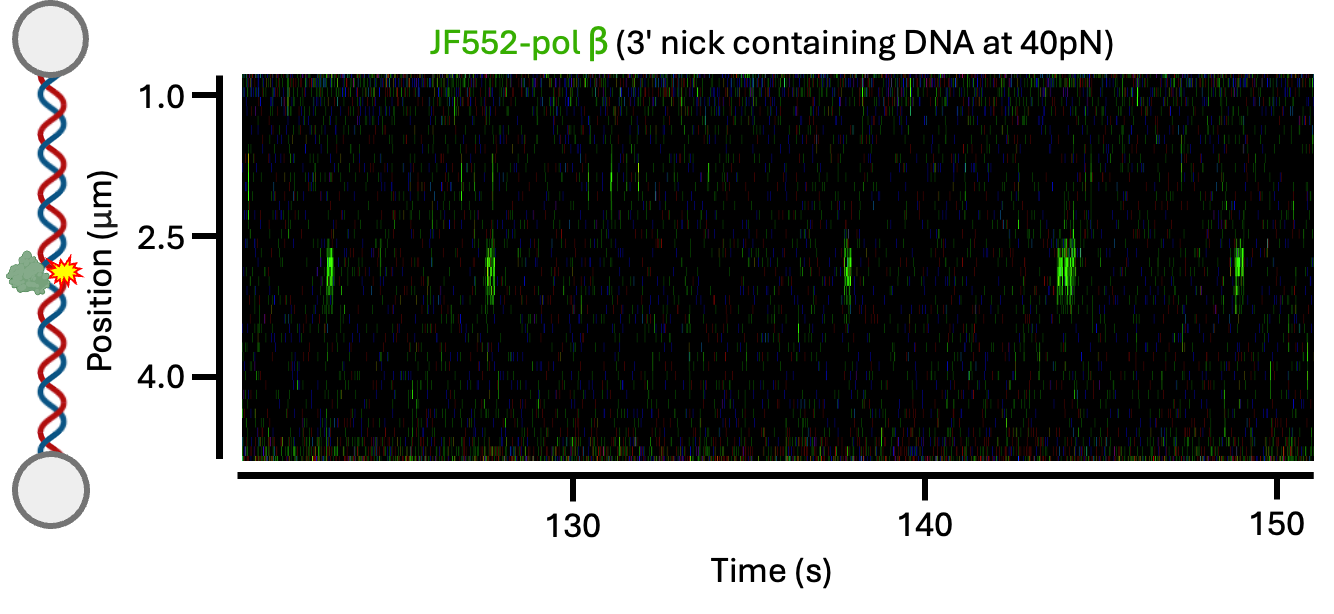

**Supplementary Figure S2. Additional kymographs of pol ꞵ on 3’ nick DNA at different forces**. (A) Example kymographs showing JF552-pol ꞵ binding to the 3’ nick stretched to 10 pN (B) JF552-pol ꞵ binding to the 3’ nick stretched to 20 pN. (C) JF552-pol ꞵ binding to the 3’ nick stretched to 40 pN.

**A**

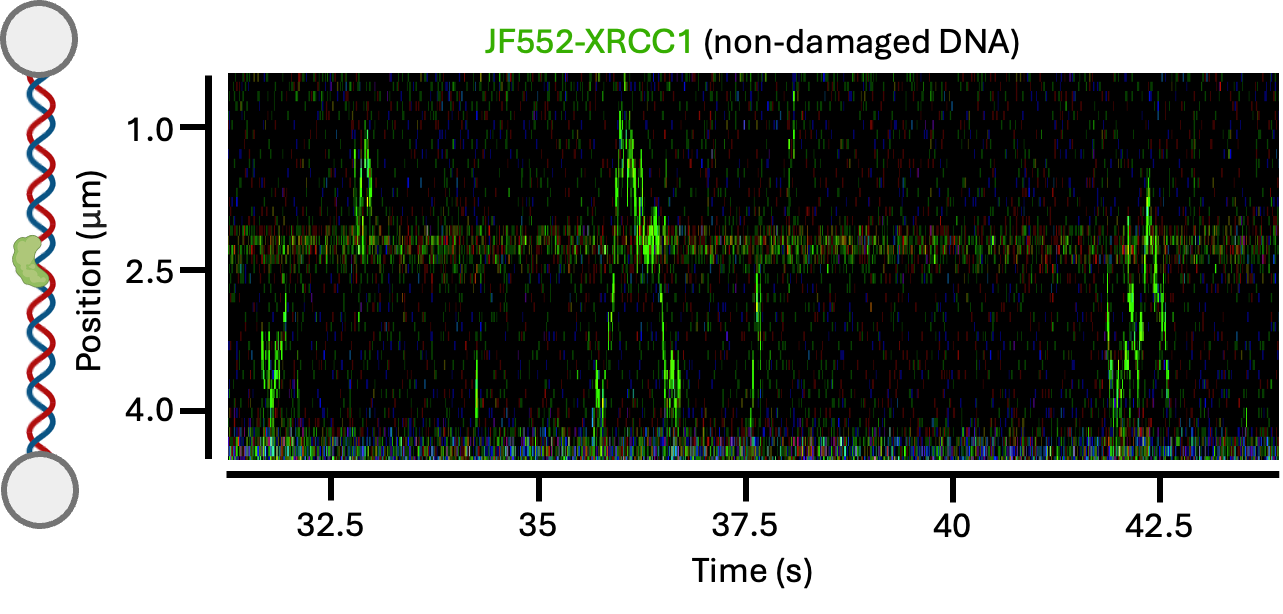

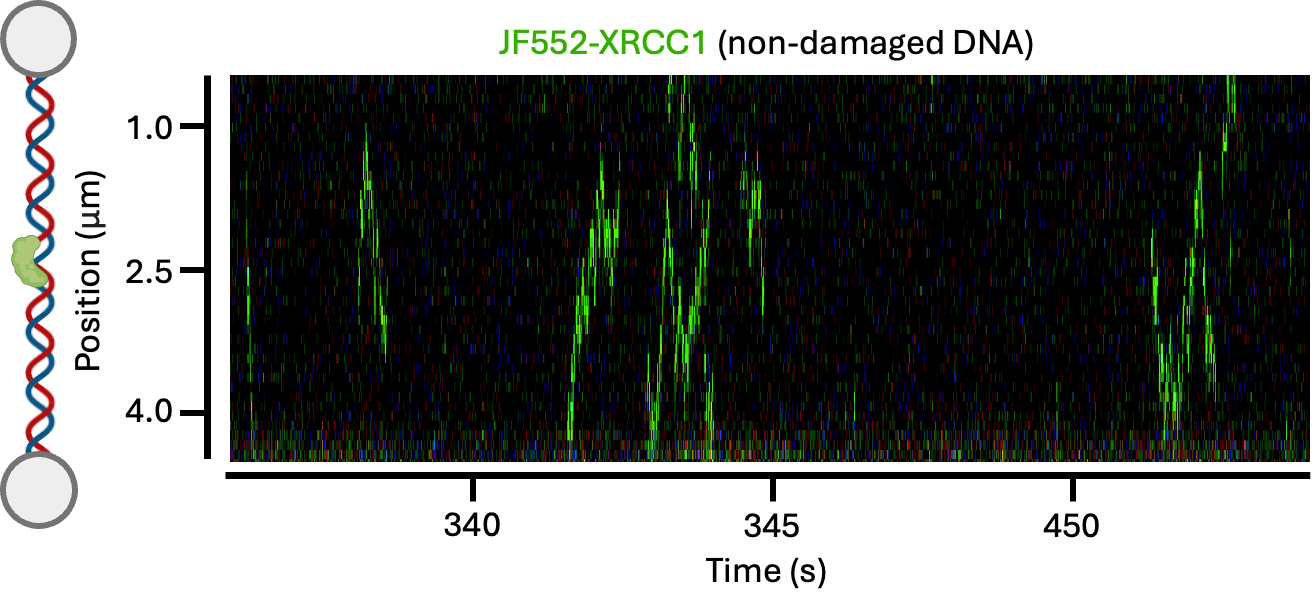

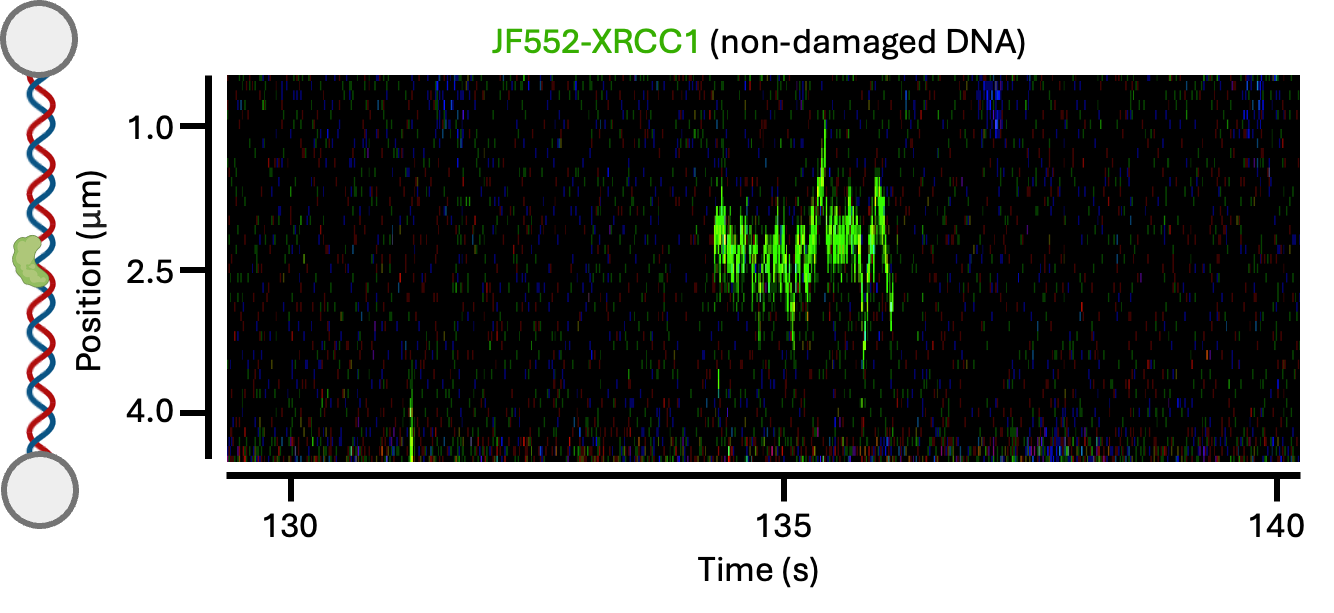

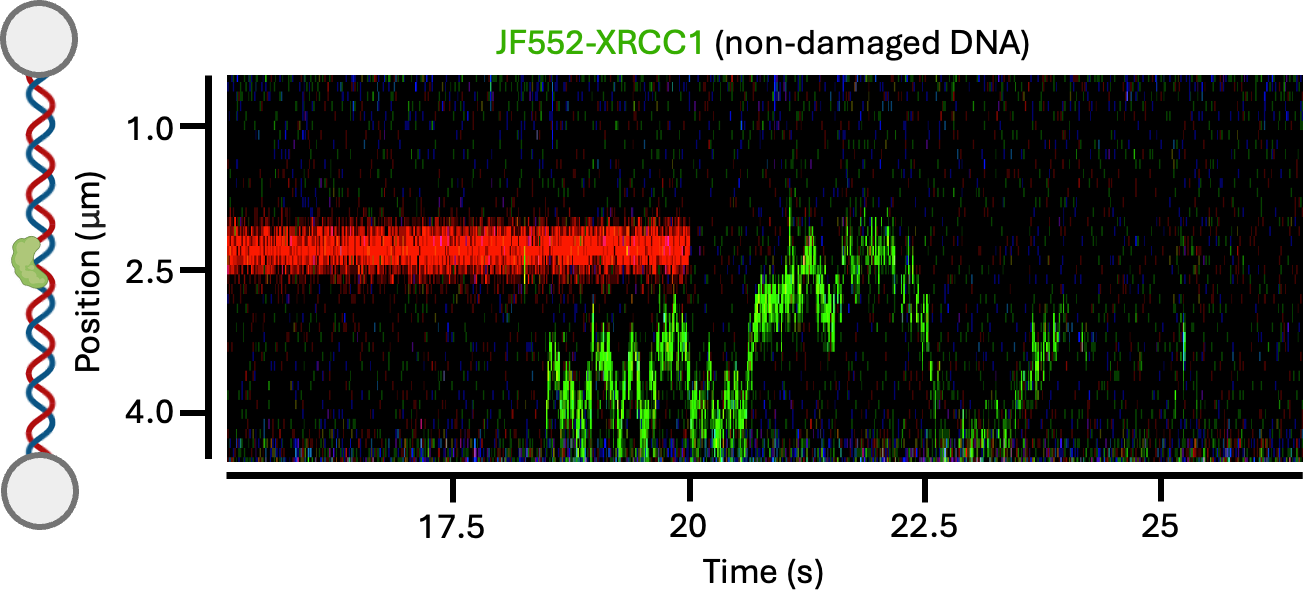

**B**

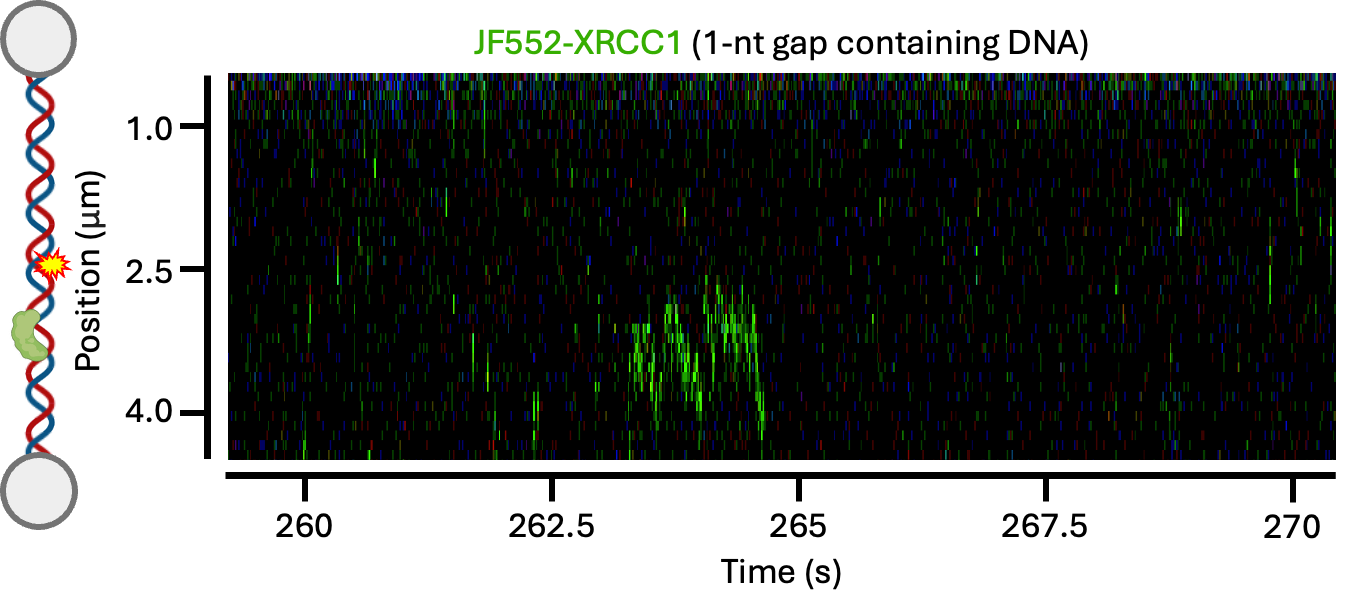

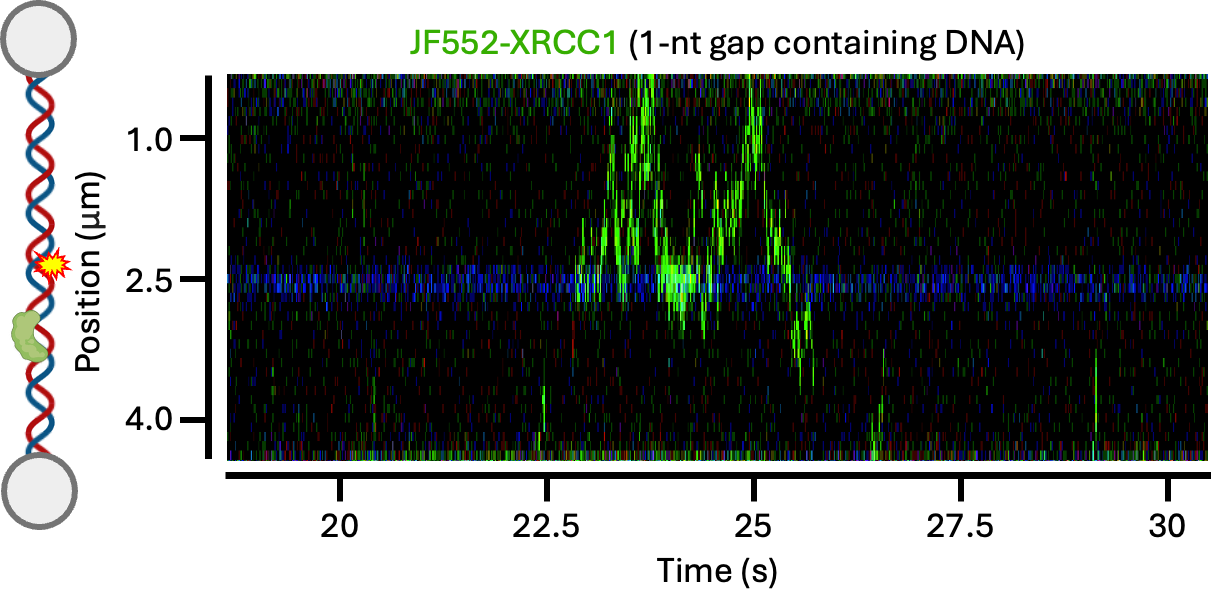

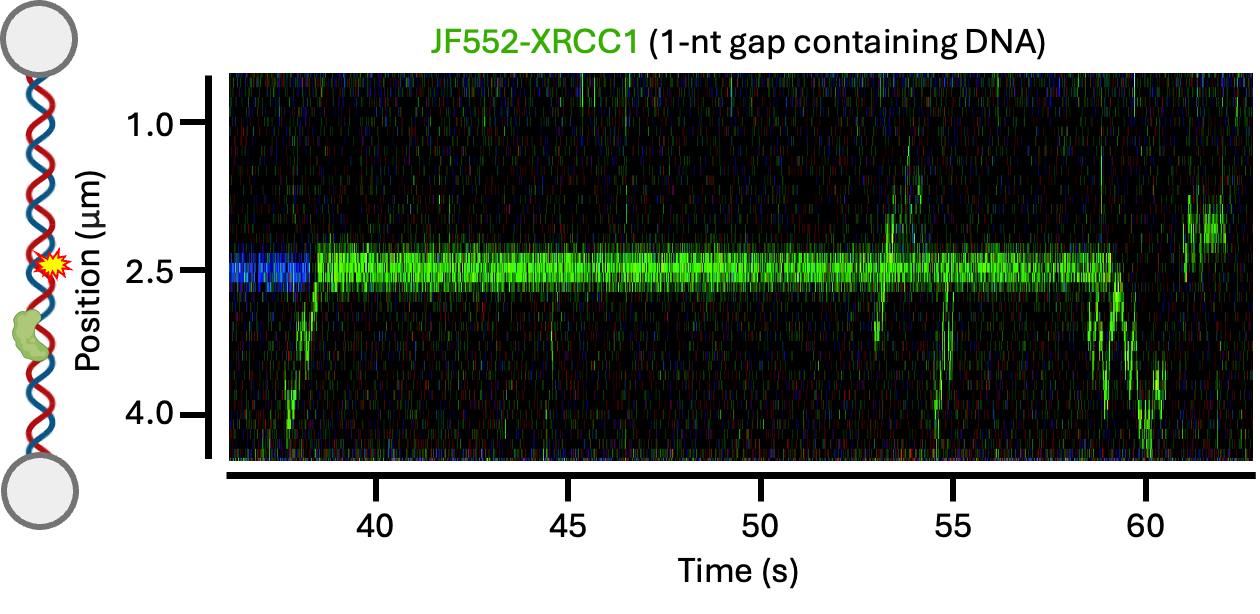

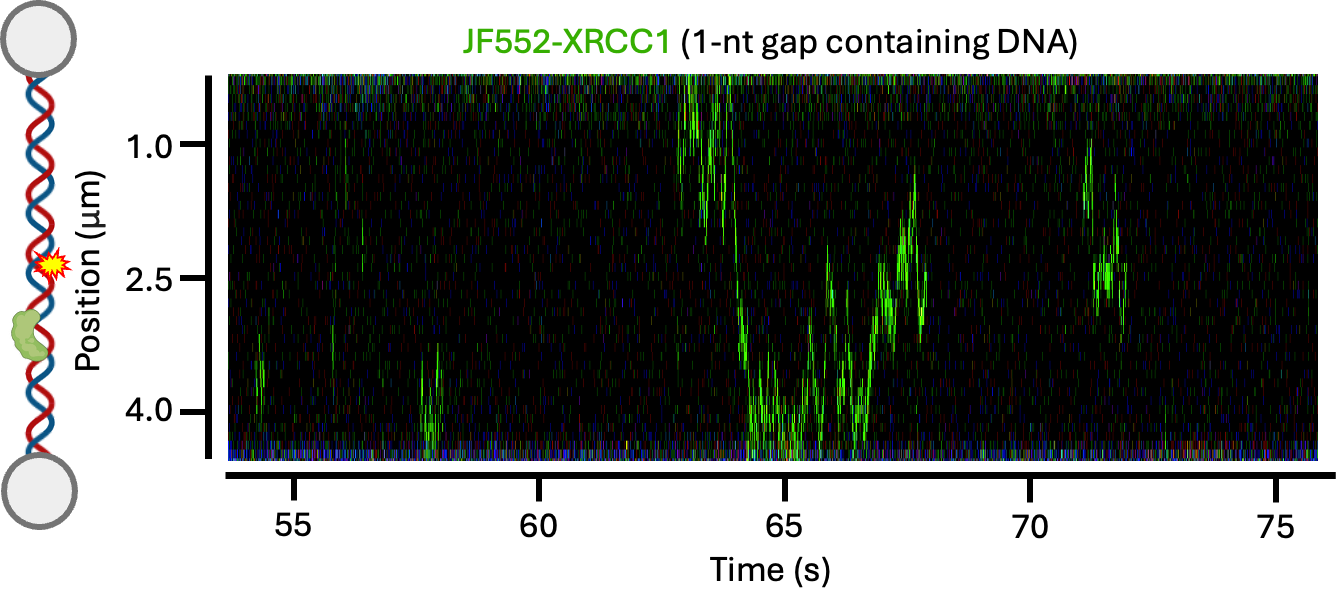

**C**

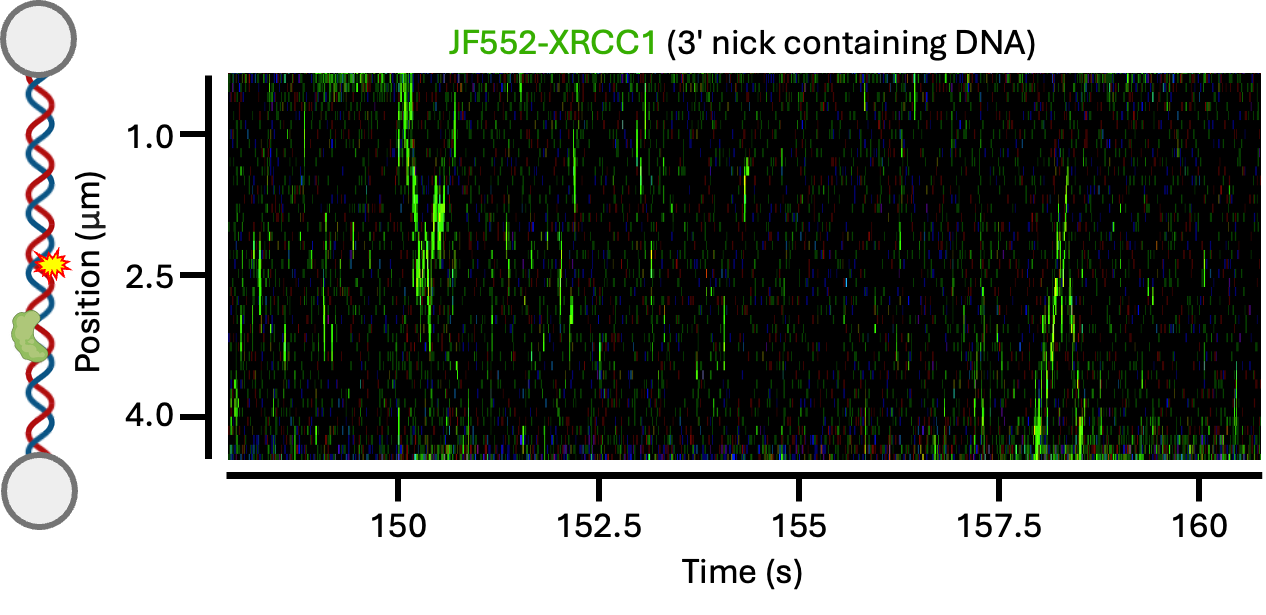

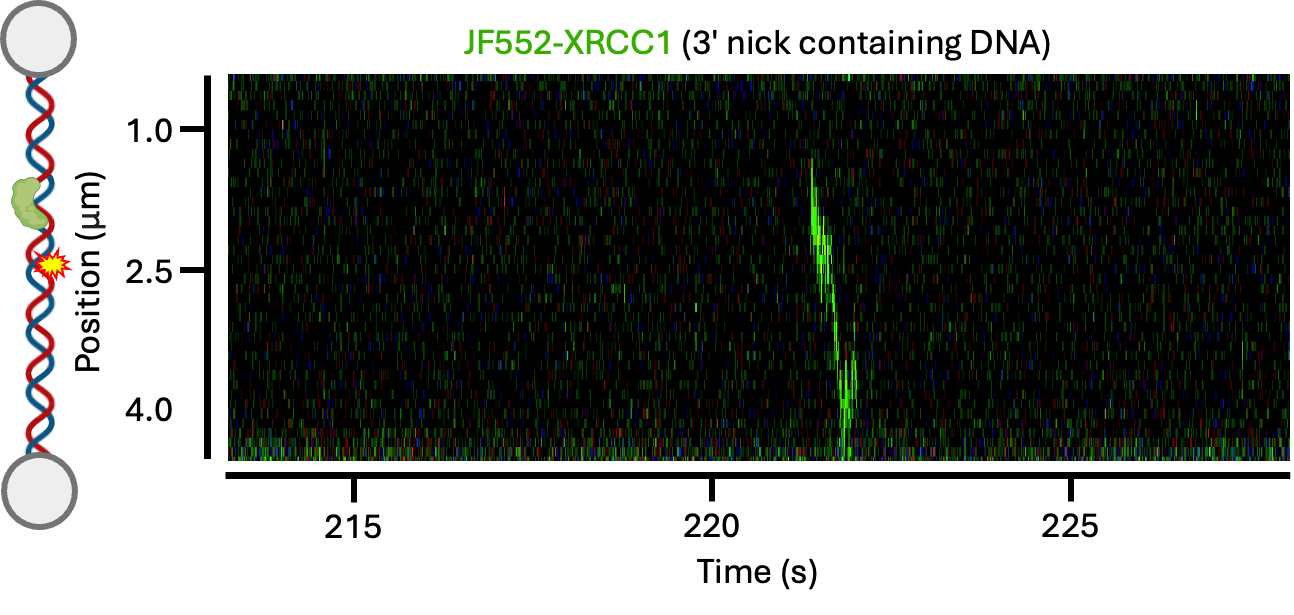

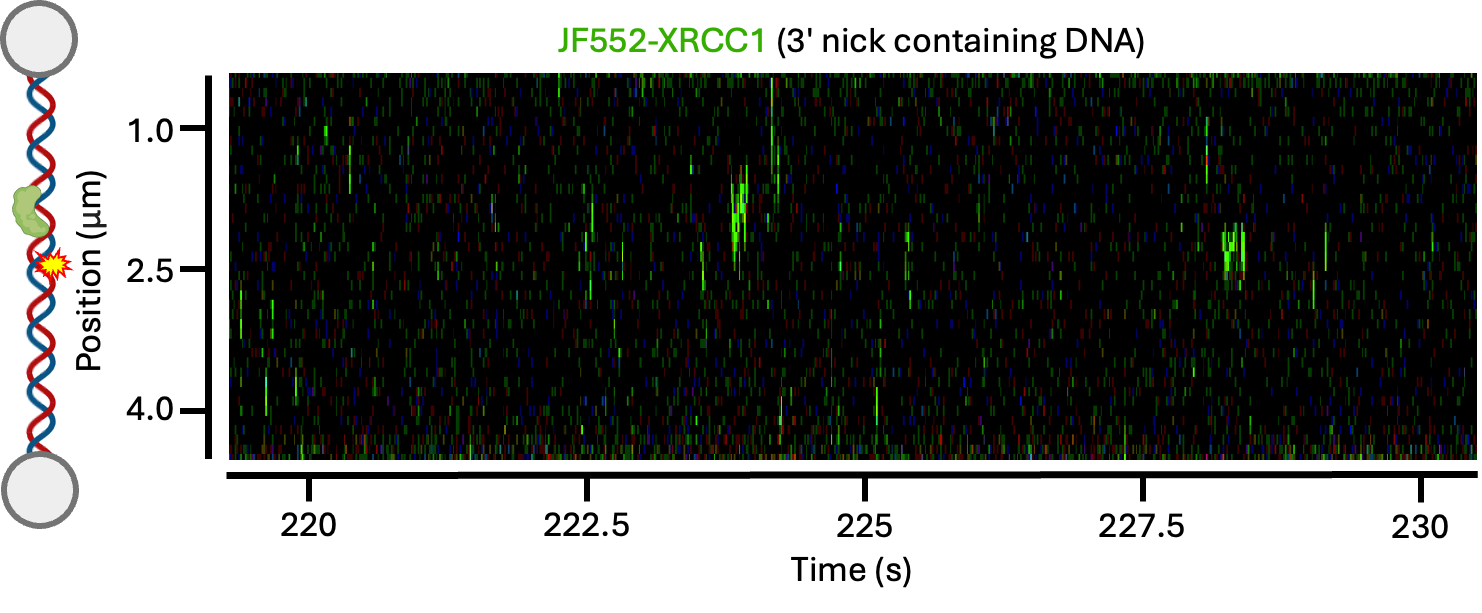

**Supplementary Figure S3. Additional kymographs of XRCC1 on different DNAs.** (A) Example kymographs showing JF552-XRCC1 moving along non-damaged DNA. (B) JF552-XRCC1 searching along DNA containing a 1-nt gap. (C) JF552-XRCC1 searching along DNA containing a 3′ nick.

**A**

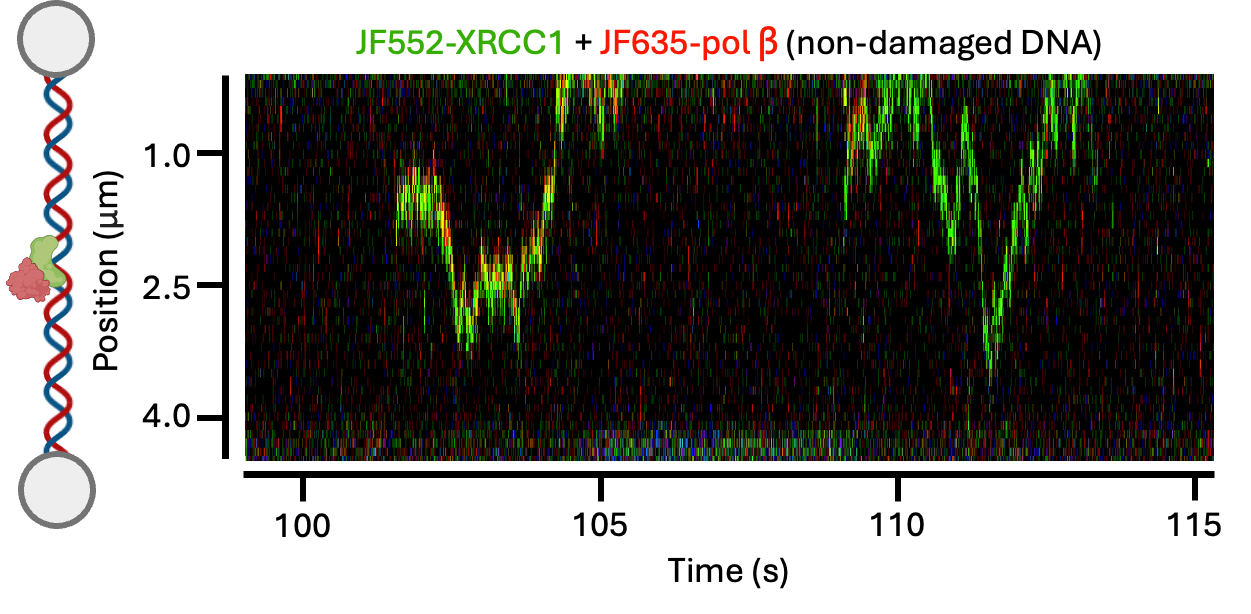

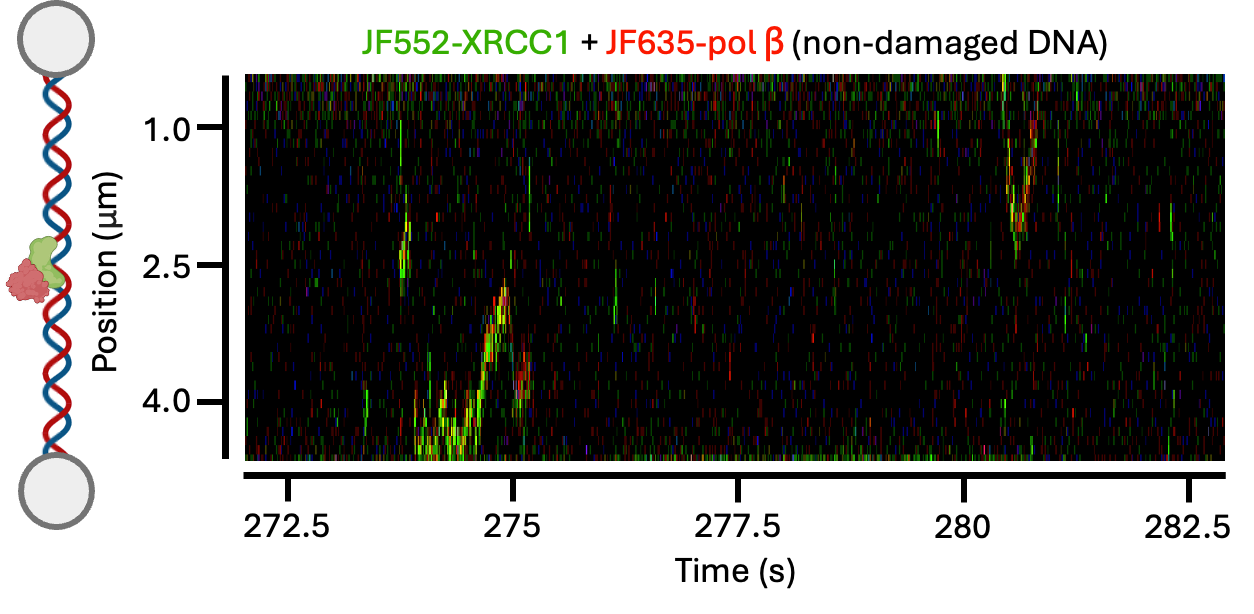

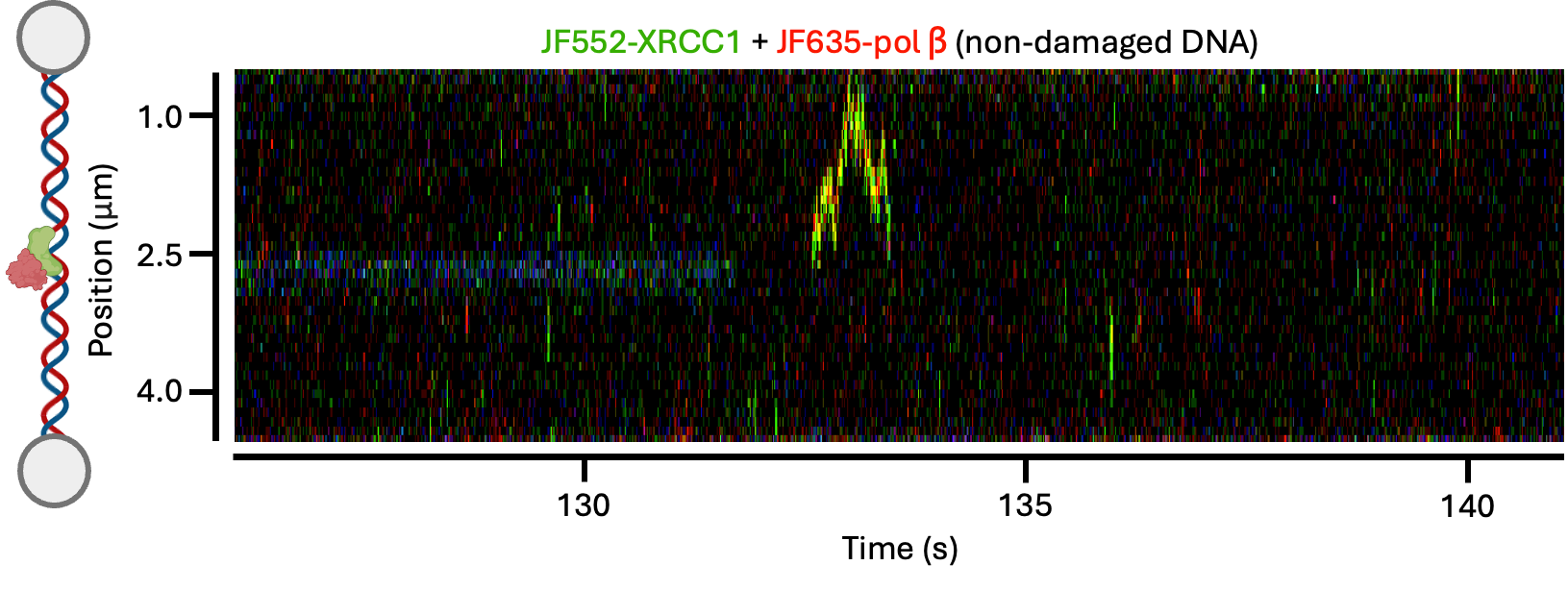

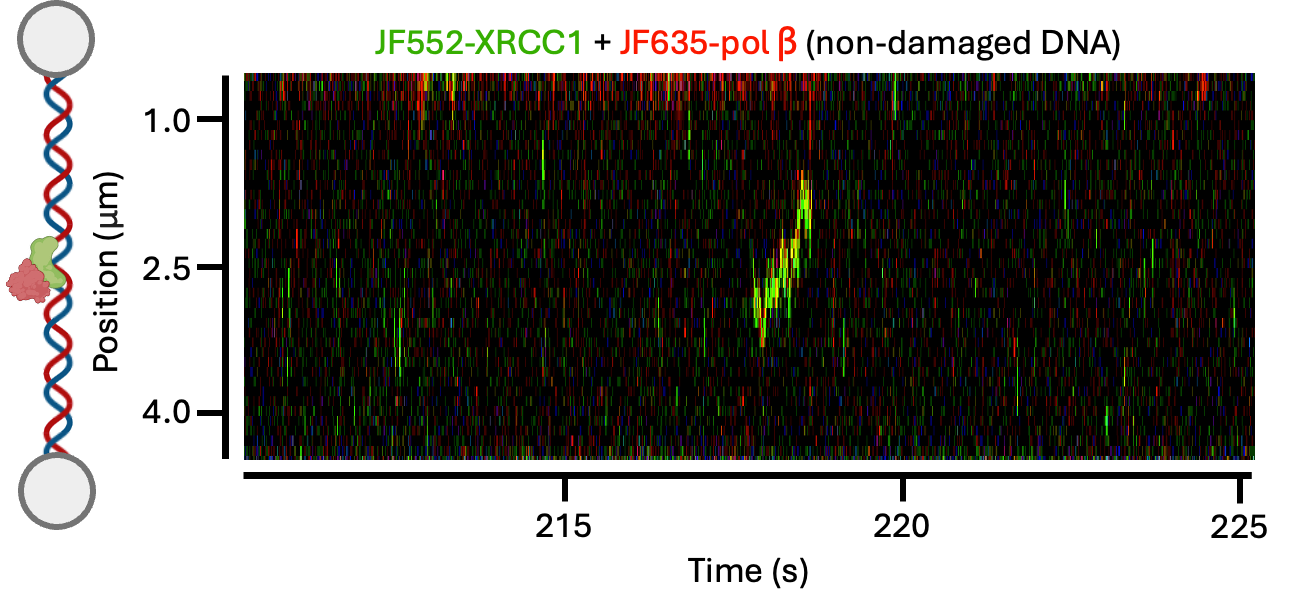

**B**

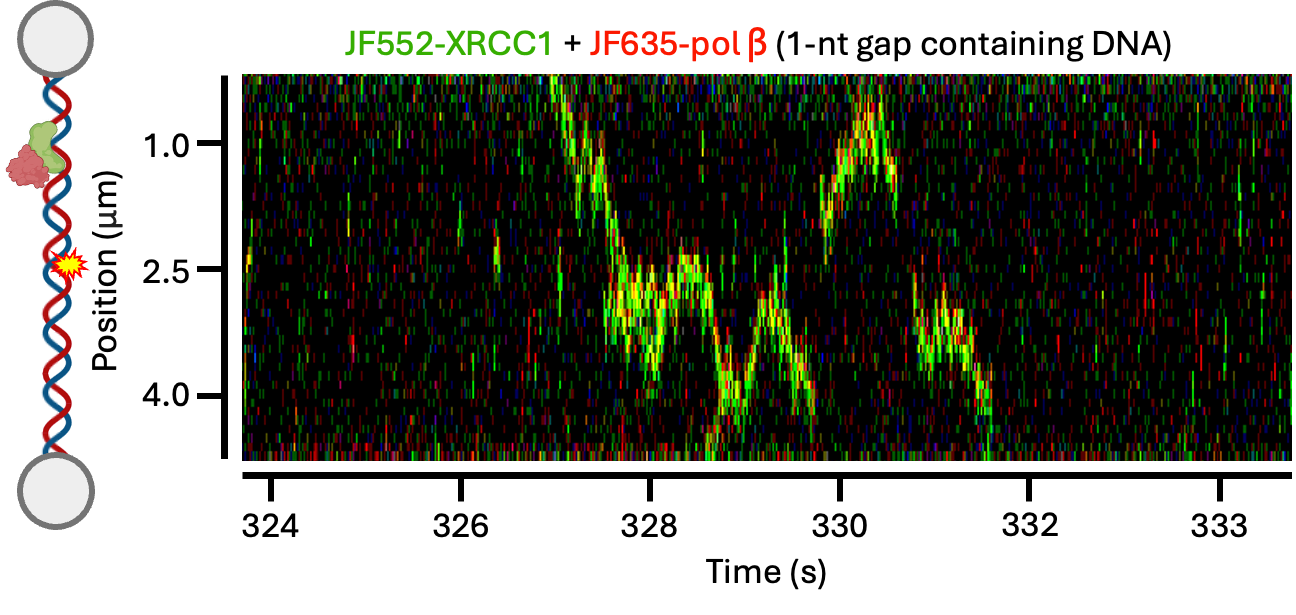

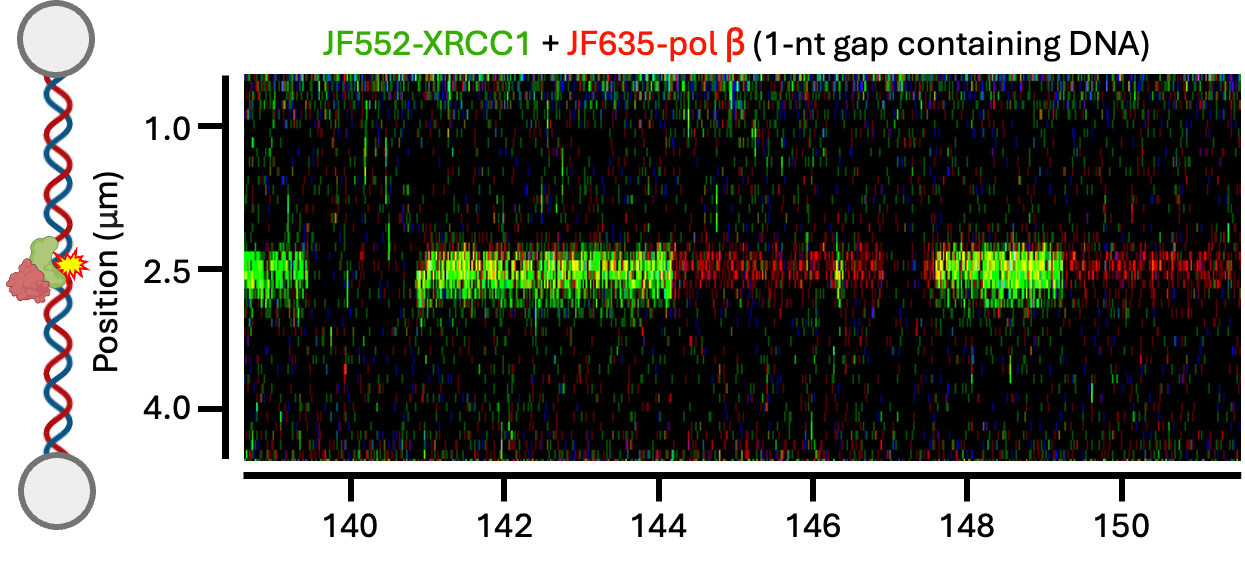

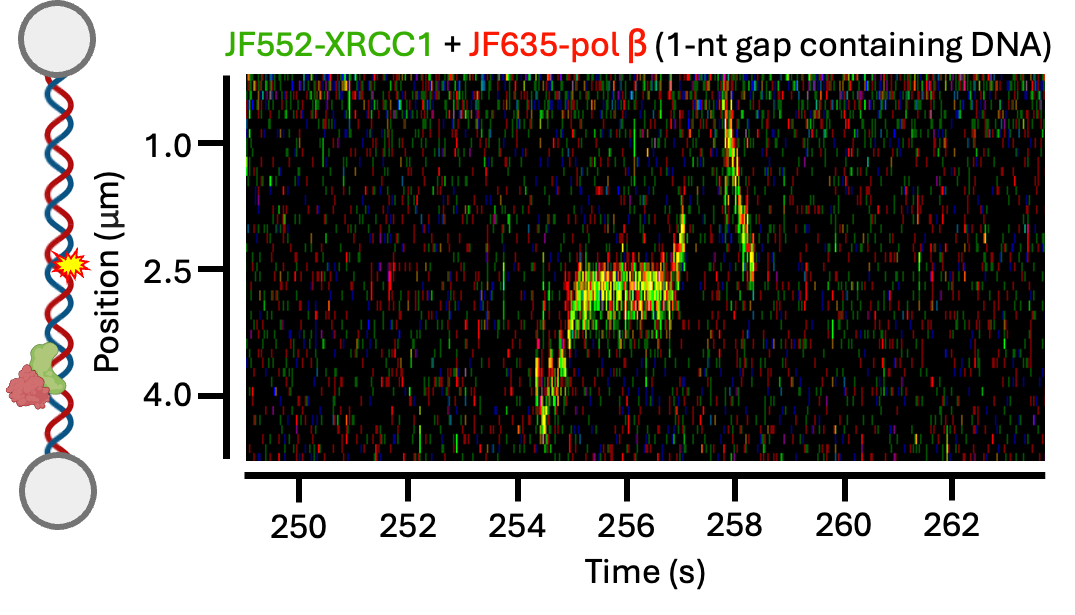

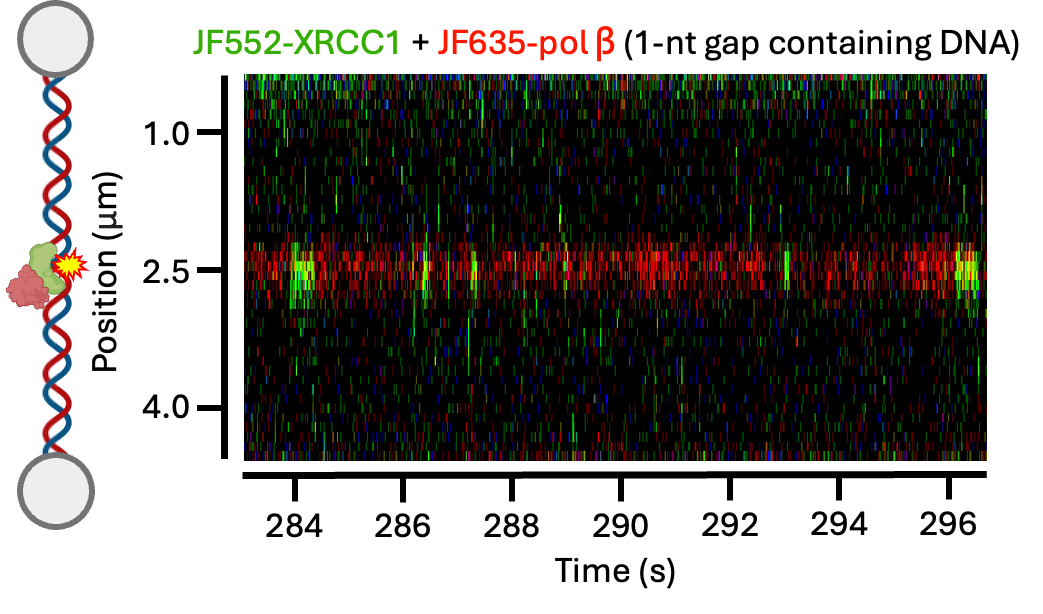

**C**

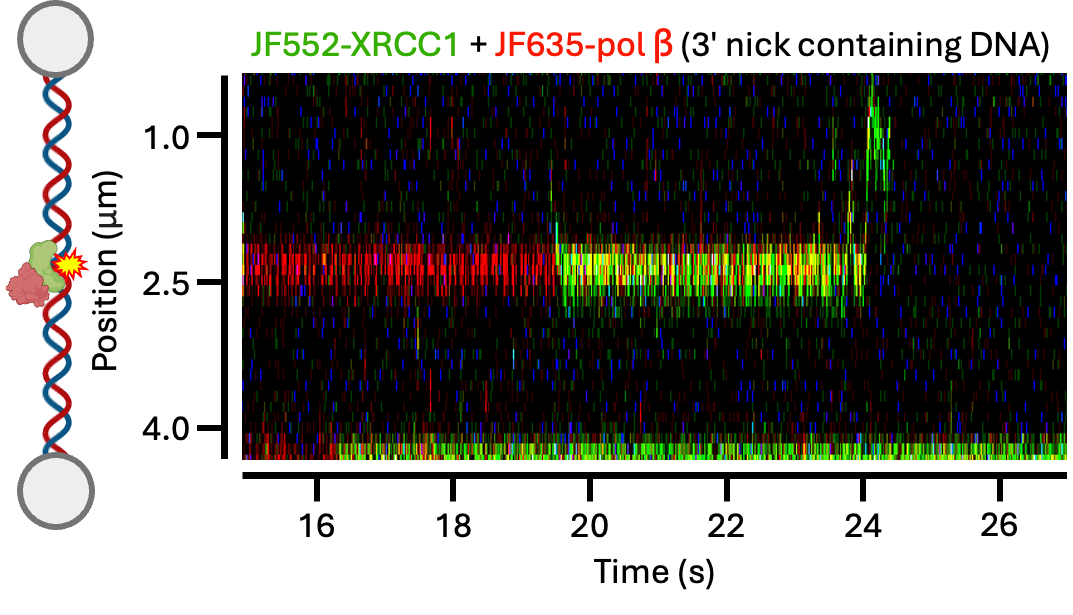

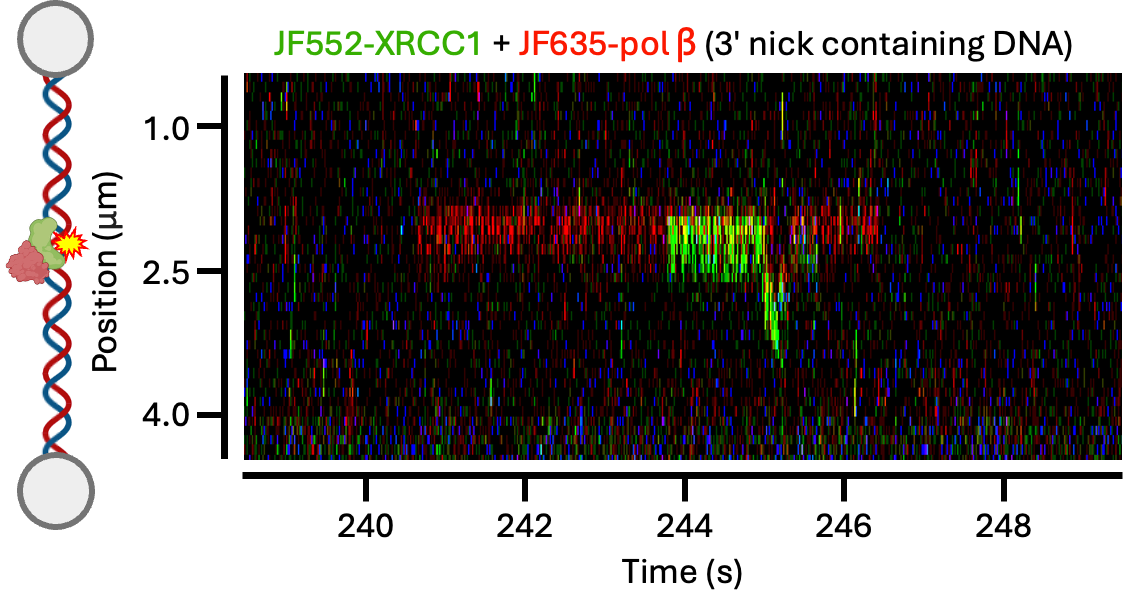

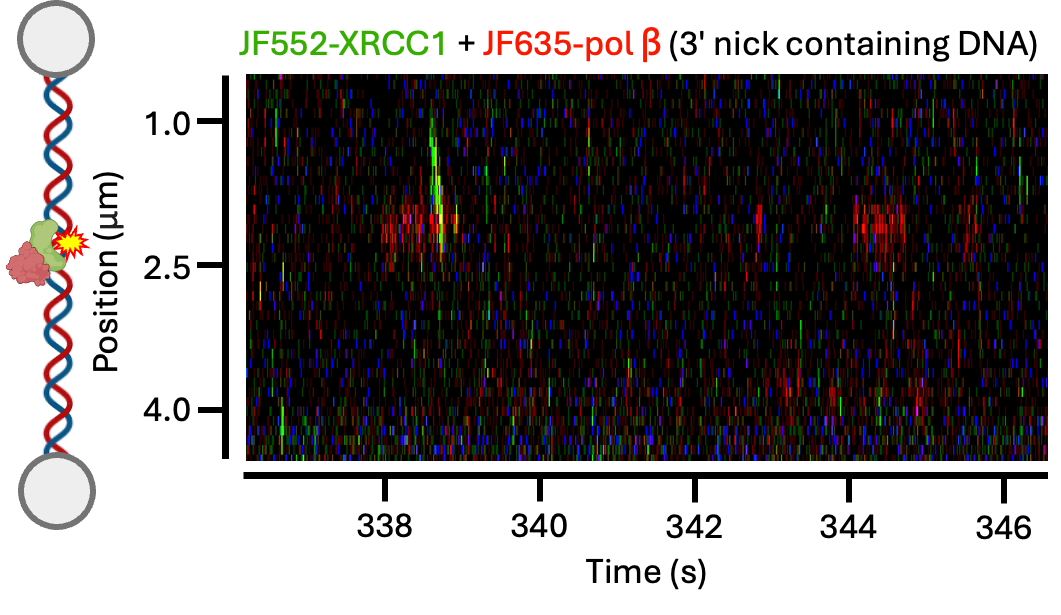

**Supplementary Figure S4. Additional kymographs of pol ꞵ-XRCC1 on different DNAs.** (A) Example kymographs showing the pol ꞵ-XRCC1 complex moving along non-damaged DNA. (B) Pol ꞵ-XRCC1 complex searching along DNA containing a 1-nt gap. (C) Pol ꞵ-XRCC1 complex searching along DNA containing a 3′ nick.

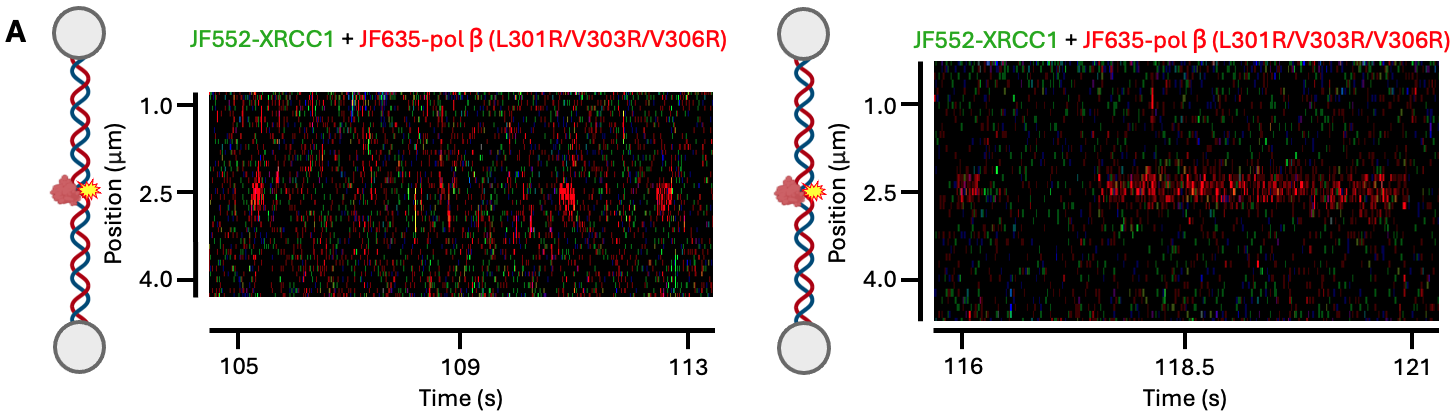

**Supplementary Figure S5. The pol ꞵ triple mutant is unable to form a motile complex with XRCC1.** (A) Pol ꞵ L301R/V303R/V306R (red) and XRCC1 (green) in the presence of a 1-nt gap. Pol ꞵ is still able to bind damage, but is unable to form a motile complex with XRCC1.

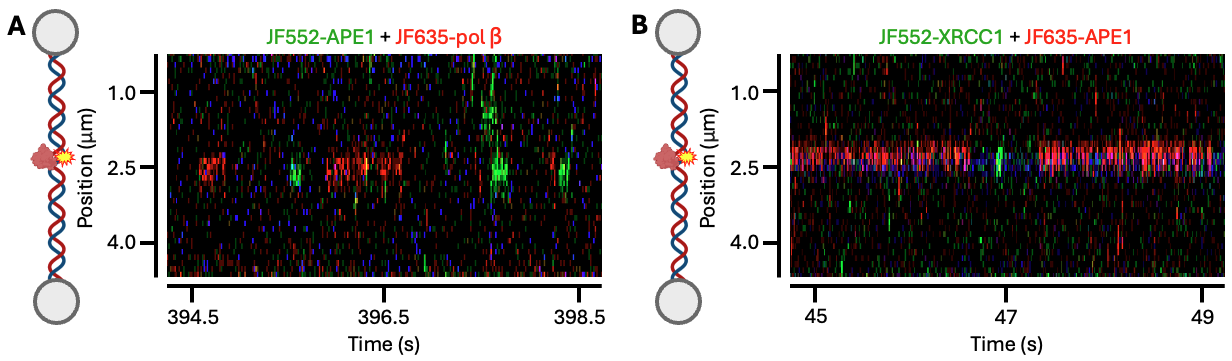

**Supplementary Figure S6. Simultaneous damage search of multiple BER proteins.** (A) Pol ꞵ (red) and APE1 (green) binding to the site of a 1-nt gap. Across 31 kymographs collected we observed 288 examples of APE1 alone binding to the 1-nt gap. (B) APE1 (red) and XRCC1 (green) binding to an AP site indicated by the Atto488 fluorophore (blue). Across 67 kymographs collected we observed 22 examples of XRCC1 alone binding to the AP site.
